# The environmental dependence of ecological interaction networks

**DOI:** 10.1101/2025.05.16.654471

**Authors:** Bastien Parisy, Alyssa R. Cirtwill, Tomas Roslin

**Affiliations:** Organismal and Evolutionary Biology Research Programme, Research Centre for Ecological Change, Faculty of Biological and Environmental Sciences, University of Helsinki; Carex EcoLogics, Bracebridge, Canada.; Department of Ecology, Swedish Agricultural University, 75007 Uppsala, Sweden; Ecosystems and Environment Research Programme, Faculty of Biological and Environmental Sciences, PO Box 65 (Viikinkaari 1), FI-00014 University of Helsinki, Finland

## Abstract

Interactions between species are often assumed to occur whenever species pairs co-occur, allowing us to reconstruct local ecological networks from a deterministic “metaweb”. Thus, a species’ set of metaweb interactions defines its Eltonian niche – the biotic conditions under which it can exist. However, both species and interactions may also have Grinnellian niches, reflecting relationships between probability of occurrence and environmental conditions. Over the past few years, we have explored how stochasticity in the occurrence of species and links, and environmental imprints on these, shape ecological interaction networks. In this perspective, we use a set of our empirically-characterised networks to show that both species and link occurrence are stochastic, with probabilities conditional on the environment. This insight improves our ability to model observed variation in network structures. We argue that future research should incorporate environmental impacts on both species and links, thus merging the Grinnellian and the Eltonian niche concept.

## Introduction

Ecological communities are conveniently described as networks, with species as nodes and interactions as links (Gravel *et al*., 2019). While community ecology to date has largely focused on the factors shaping local species richness and composition, the interactions between these species forms another facet of community structure. This was recognised early on by the father of modern animal ecology, Charles Elton (Elton, 1927). While the foundational Grinnellian concept of species niches focused on the environmental conditions within which a species maintains positive population growth, Elton defined the niche as the set of interactions between a focal species and other taxa in its environment (Chase & Leibold, 2003). In this paper, we explore the links between the Grinnellian and the Eltonian niche concepts – and call for an urgent synthesis between the two (Gravel *et al*., 2019; Cirtwill *et al*., 2019).

The Eltonian niche has recently experienced a revival in the context of ecosystem functions and ecosystem services (Moll *et al*., 2021; Curras *et al*., 2025). In essence, species interactions form the underpinnings of ecosystem functions (Wirta *et al*., 2021, 2022; Roslin, 2024). Interest in network ecology has revolved around – among other things – the factors shaping the relative stability of networks, the resilience of networks to species loss, the topology of networks, and the dynamics of networks of different structure (Mace *et al*., 2018; Lu *et al*., 2020; Heading & Zahidi, 2023; Naeem *et al*., 2016). In contrast, the link between global and community change has been built on the Grinnellian niche concept, which offers a direct connection between changes in environmental conditions and changes in species distributions. As a result, environmental impacts on species interactions have been less studied (Pellissier *et al*., 2018) and the field of community change has been somewhat detached from the field of changes in ecological networks (Olesen *et al*., 2010; Woodward *et al*., 2010). Yet, both perspectives on the species niche are now more topical than ever, and a synthesis between them is sorely needed (Gravel *et al*., 2019; Cirtwill *et al*., 2019).

From a network perspective, the Eltonian niche concept can be defined as the set of species with which the species interacts (Gravel *et al*., 2019). While the concept may seem seductively simple, it becomes more complex when applied to a community consisting of tens to tens of thousands of species and their interactions. Empirical studies of a species’ biotic setting (i.e., set of co-occurring species) record the interactions occurring at a particular time and place (i.e., a point within a Grinnellian niche space). These records are commonly pooled across times and places to create a “metaweb”, which purports to describe the biotic context of each species within it – but neglects environmental influences entirely (Fig. 1). Moreover, metawebs frequently combine information from various sources, such as literature records of species interactions, species association patterns across motley samples, or expert opinion (sometimes validated through experimentation) (Roslin & Majaneva, 2016). These disparate sources may include varying amounts of information about species’ Grinnellian niches, but combining them in a metaweb such as the classical food webs of the Chesapeake Bay (Baird & Ulanowicz, 1989), Benguela current (Yodzis, 1998), and the island of St. Martin (Goldwasser & Roughgarden, 1993) obscures any such insights.

**Figure 1:**
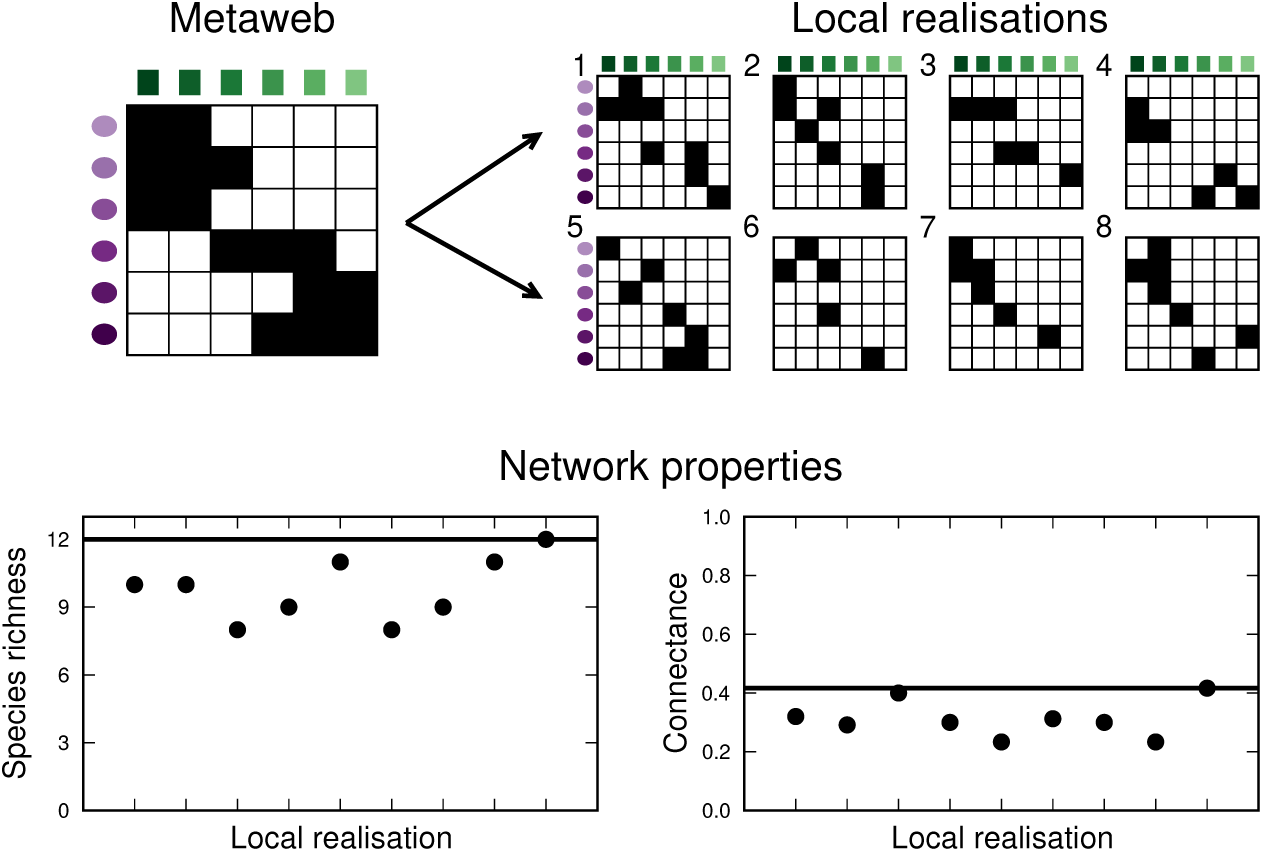
For a given set of species and feasible interactions between them (“metaweb”, top left), only some species and interactions will occur at a given space or time. This results in a variety of local realisations (top right) of the network. Each realisation of the network will have network properties, such as species richness or connectance (dots in bottom panels), that may differ from the metaweb values (black lines in each panel).

A synthesis between the Grinnellian and the Eltonian niches is urgently needed to address topical questions about how communities of species will respond to ongoing ecosystem change. First, we may ask how the role of individual species will change with a change in their environment (Cardoso *et al*., 2011; Peterson, 2011; Rosado *et al*., 2016). Here, the network literature has focused on the impact of invasive species anthropogenically introduced to new regions (Memmott & Waser, 2002; Albrecht *et al*., 2014; Traveset *et al*., 2016), but less on the general rebuild of ecological interaction networks as a consequence of climate change (but see Tylianakis *et al*., 2008; Bascompte & Stouffer, 2009; Fontúrbel *et al*., 2021. Second, we may ask how community structure will change with global change (Murphy & Romanuk, 2014; Newbold *et al*., 2015; Komatsu *et al*., 2019). In this context, the assumption that species will change independently of their food resources and enemies is necessarily na ï ve (Endara & Coley, 2011; Mougi, 2022). Third, we may ask how ecosystem functions and services will change with global change (Bongaarts, 2019; Begon & Townsend, 2021; Fiedler *et al*., 2021; IPCC, 2023). To appreciate this, we should consider the sum of partly counterbalancing species interactions, not interactions one by one.

Moreover, any investigation of changing ecosystem functions and services should consider the possibility that interactions between species pairs may *also* have a Grinnellian niche. In other words, a given link between two species may occur with a different probability under different environmental conditions (Poisot *et al*., 2012, 2015), adding to variation in the Eltonian niche of the species with variation in environmental conditions (Parisy *et al*., 2024a,b). Common reasons for environmental forcing of interaction structure include physiologically determined responses to temperature and phenological shifts among interacting trophic layers. Among the latter, effects of changing environmental conditions differ depending on species’ trophic levels (Schmidt *et al*., 2016, 2023; Roslin *et al*., 2021; Bell-Dereske *et al*., 2023). In the context of climate change, changes in environmental conditions may also cause shifts in competitive constellations within layers (Su *et al*., 2024; Wootton *et al*., 2024; Tiusanen *et al*., 2020). These shifts may sometimes be mediated by links to other trophic layers, such as competition among plants for access to pollinators or *vice versa* (Tiusanen *et al*., 2020; Redr *et al*., 2025). We note that species may also switch interaction partners in response to changing abundances, even if their earlier interaction partners are still present and active (Carnicer *et al*., 2009; Poisot *et al*., 2015). Just like changes in phenology, this preference switching or interaction rewiring would result in changes to species’ direct and indirect interactions (i.e. their roles in the network) over time, while these changes would not be associated with species turnover. Interaction networks are therefore highly conditional on the environment rather than constant (Fig. 2; Gravel *et al*., 2019).

**Figure 2:**
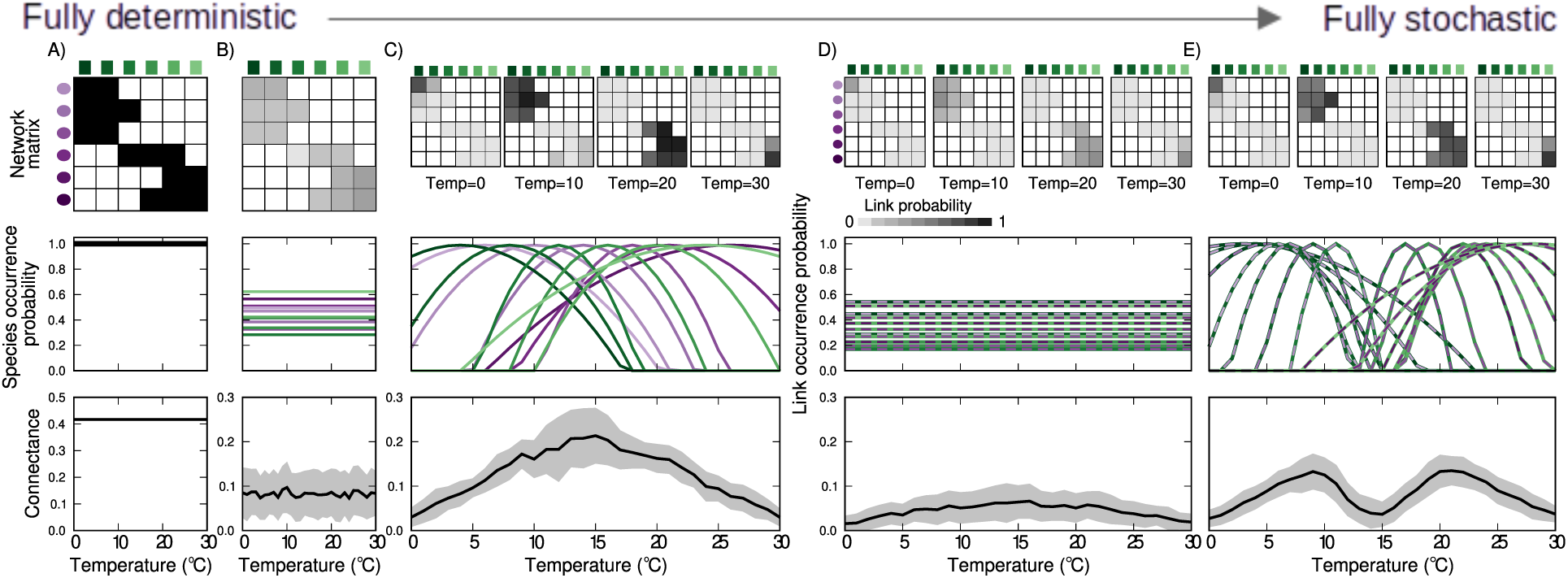
The stochastic vs deterministic view on species interactions. Columns of panels show impacts on different aspects of network structure as we horizontally move from a deterministic view to one which incorporates Grinnellian niches of species and their interactions. The top row shows the network matrix, in which cell shading indicates the probability that a link occurs, the middle row shows the probability of each individual species (A-C) or link (D-E) occurring at each temperature, and the bottom row shows the mean (±SD) connectance of 100 simulated observed networks for each temperature. (Note change in scale of the connectance axis between A and B–E.) Impacts are compared among six scenarios: (A) Species occur with probability 1 at all temperatures, and feasible interactions always occur. (B) Species occur with a constant probability, which differs between species. This reduces the probability that interacting species will co-occur and interact, leading to lower, and variable, connectance. (C) Species have unique Grinnellian niches with respect to temperature (curves in middle panel), leading to varying probabilities of co-occurrence. As in B), species always interact when they co-occur with a feasible partner. Columns D–E combine these Grinnellian niches with interaction stochasticity: (D) Interactions are stochastic with constant probability across temperatures, lowering the probability of any interaction occurring and decreasing the connectance of the resulting observed networks. (E) Some interactions are more likely at cooler temperatures, others at warmer temperatures (depending upon the temperature optima of the species involved). In this example, this results in somewhat higher connectance at moderately low (5-10^◦^C) and moderately high (20-25^◦^C) temperatures than when interaction probabilities are constant across temperatures (D). Note that the probabilities of link occurrence in D are the average values of the curves in E; likewise, the probabilities of species occurrence in B are the average values of the curves in C. For simulation methods, see Appendix *S1*.

While the idea that ecological network structure depends on environmental conditions may sound like a truism, its consequences have been neglected. To understand changes in community composition, we as a field have generally modelled species as responding one-by-one to changes in their abiotic environment, as captured by the Grinnellian niche (Elith & Leathwick, 2009; Antão *et al*., 2022). At the same time, we have stuck to the metaweb approach, assuming no environmental impacts on the Eltonian niche of the species (Rosado *et al*., 2016) and that local species interactions can be deterministically inferred from species co-occurrences (Schlegl *et al*., 2014; Ho *et al*., 2022; Mendes *et al*., 2024).

More recently, some authors have argued for a probabilistic view of species interactions, emphasising that pairs of species will not always interact when they encounter each other (Cirtwill *et al*., 2019; Gravel *et al*., 2019; Poisot *et al*., 2016). Even a stochastic view of interactions, however, neglects the possibility of environmental influences on the local probability of an interaction occurring – the fullest integration of Eltonian and Grinnellian niches (Fig. 2, Gravel *et al*., 2019). When environmental and interaction data are collected in tandem and in quantity, it is possible to assess how probabilities of species and interaction occurrence vary with environmental variables such as temperature (i.e., to estimate their Grinnellian niches) and to incorporate these estimates into network models.

In this Perspective, we systematically demonstrate the imprints of stochasticity and environmental dependence on ecological interaction networks. To this end, we first use a series of scenario simulations to explore how network properties will vary with our explicit assumptions regarding the nature of species and link occurrence. We then draw on empirical data from a large set of well-characterised networks to ask whether interactions within these networks are stochastic or deterministic in nature and whether basic interaction probabilities vary with environmental conditions, i.e. whether links can be characterised by a Grinnellian niche. Having found that they can, we predict the structure of the observed networks including different aspects of stochasticity or environmental influence to identify the approach that best captures differences in observed network structure.

In all analyses, we describe interaction networks at different scales, providing different insights into each species’ biotic setting: we consider macro-scale or whole-network measures (connectance, nestedness) as well as meso-scale measures (network motif profiles and species motif roles (Baker *et al*., 2015); Fig. 3). While macro-scale measures provide an efficient summary of network structure, they can mask substantial fine-scale variation (Stouffer *et al*., 2012; Baker *et al*., 2015; Simmons *et al*., 2019a). Both the motif profile of a network (counts of unique patterns of interacting species (Milo *et al*., 2002, 2004; Stouffer *et al*., 2007)) and species’ motif roles (frequencies of species’ appearance of each position in each motif) capture more of this fine-scale structure, while also describing direct and nearby indirect interactions between species – i.e., interactions within a few ‘degrees of separation’ (Cirtwill & Stouffer, 2015). This focus on nearby indirect interactions is ecologically relevant, as it is generally expected that species will have the strongest effects on their direct interaction partners and progressively weaker effects on species to which they are only indirectly connected through long paths (Cirtwill *et al*., 2018). Smaller motifs are therefore likely to capture most of a species’ strong interactions, while larger motifs will include progressively weaker indirect interactions. Here, we consider motifs of two to six species.

**Figure 3:**
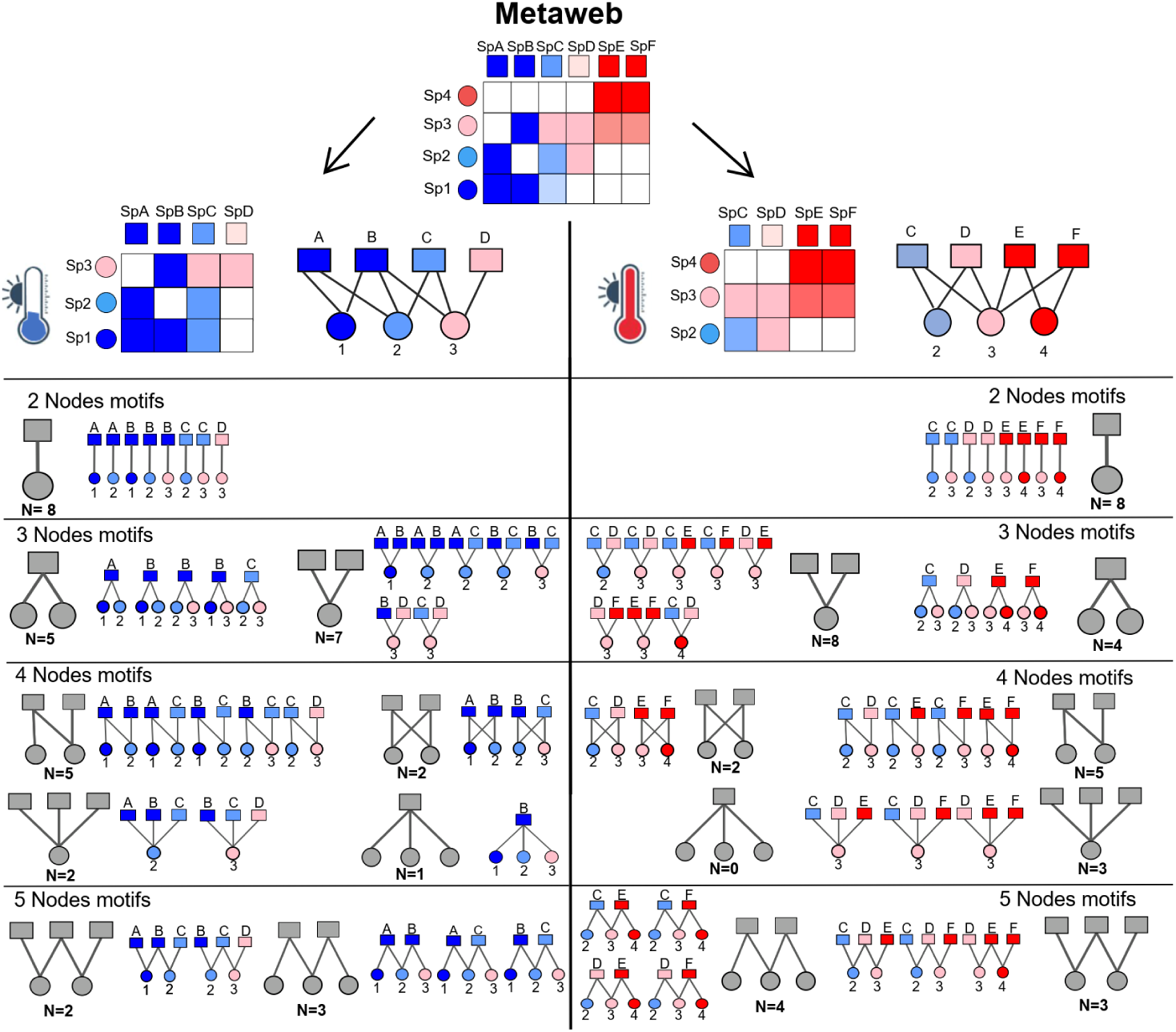
Describing a network’s motif profile can capture meso-scale changes in network structure which are masked when focusing on macro-scale measures like connectance. Here, local realisations of a bipartite network of consumers (top, circles) and resources (bottom, squares) differ slightly between cold (left) and warm (right) sites. Specifically, resource A is present at the cold site only and resource D is present at the warm site only. Note that, as numbers of species and links are identical between sites, connectance is constant in this example. However, due to the change in *identity* of the species present, the motif profile of the network has changed. To calculate the motif profile of a network, the network is first decomposed into all unique sets of *n* species and *all* of their interactions. If this sub-network is a single compartment (i.e., all species are directly or indirectly connected to all others), it is then matched to one of the possible motifs of size *n* as shown below. Repeating this process gives a count of how many times each motif of size *n* appears in the network. As a compromise between describing network structure in detail and computational limitations, bipartite networks are typically described using motifs of size 2-6. Note that a group of *n* species is only assigned to a motif of size *n* and not used to identify smaller motifs. The motif profile of a network (vectors in bottom shown for motifs of size 2-4) is simply a vector of the counts of each motif found in each network.

## Materials and methods

### Methods overview

To comprehensively assess environmental imprints on ecological interaction networks, we combine analyses of empirical and simulated data. Using a toy network, we first illustrate the multiple ways in which stochastic events and environmental dependencies are likely to impact network structure (Fig. 2, Appendix *S1*). Turning to empirical data, we systematically probe for imprints of stochasticity and environmental context-dependence. We first test whether interactions within these networks are stochastic or deterministic in nature and whether basic interaction probabilities vary with environmental conditions. We then simulate networks under different scenarios of environmental impacts, demonstrating drastic differences in change along environmental gradients. In all analyses, we resolve imprints on different scales: network-level summary properties (number of links, connectance, and nestedness) and meso-scale network motif profiles and species motif roles (Fig. 3. In all analyses, connectance and nestedness (specifically, NODF) are calculated using the ‘networklevel’ function in the R (R Core Team, 2024) package *bipartite* (Dormann *et al*., 2008, 2009). Following Baker *et al*. (2015), we define a network’s motif profile as the count of all motifs containing two to six species. Each motif is a unique arrangement of *n* species and all interactions between them (Fig. 3; Milo *et al*., 2004) and each therefore represents a unique set of direct and indirect interactions among species (Stouffer *et al*., 2007; Simmons *et al*., 2019a). Motif profiles are calculated using the R (R Core Team, 2024) function ‘mcount’ in the *bmotif* package (Simmons *et al*., 2019b). Species motif roles are counts of the number of times each species appears in each *unique position* in each motif (Stouffer *et al*., 2012; Baker *et al*., 2015). To focus on differences in which motifs a species participates in, rather than the total number of motifs, we convert the vector of counts of each position into frequencies by dividing all counts by the total number of times the species appears in any position in any motif. Note that some studies (e.g., Simmons *et al*. (2019a)) normalise motif frequencies within each size class rather than across sizes, but we wish to capture changes in the relative proportions of larger and smaller motifs in a species’ role. Motif roles are calculated using the function ‘node positions’ in the R (R Core Team, 2024) package *bmotif* (Simmons *et al*., 2019b).

### Empirical data

As a basis for analyses of empirical data sets, we use the two largest sets of systematically collected data on ecological interaction networks in the Arctic of which we are aware. To resolve variation in time, we use data on plant—pollinator interactions from Zackenberg, Greenland, collected during five different years (1996, 1997, 2010, 2011, and 2016; Olesen *et al*., 2008; Rasmussen *et al*., 2013; Cirtwill *et al*., 2023). To compile these networks, two focal individuals of each plant species present were observed for 40 minutes per day and all insects visiting open flowers were collected (Appendix *S2*). These observations were repeated each day with fine weather during the entire snow-free season (see Olesen *et al*., 2008; Rasmussen *et al*., 2013 for full details). In 2016, pollen carried on insects’ bodies was sampled as well as flower visitors (Cirtwill *et al*., 2023). To maintain consistency across years, we use only the flower-visitor data from 2016. The Zackenberg metaweb includes 39 plants, 115 flower-visiting insects, and 799 interactions. This data set is henceforth referred to as the “pollination network”.

To resolve variation in space, we use data on plant—fungus interactions collected across the Arctic during the summers of 2020 and 2021 (Parisy *et al*., 2024a,b). This dataset has been generated by a coordinated sampling of roots from 12 widely-distributed plant taxa across the global Arctic (Appendix *S2*). Specifically, seven sites were selected across a gradient of 14.5^◦^ latitude. Within each site, networks of plants and root-associated fungi were characterized in 8-35 plots (c. 25m^2^). Each plot contained three to five of the 12 target plant species; root-associated fungi were identified for each plant using DNA metabarcoding while the fungal community present in adjacent soils was characterised using eDNA. In total, the networks at each site included 149-369 fungus species and 365-1167 unique interactions. This data set is henceforth referred to as the “plant-fungus network”.

### Are interactions in empirical networks deterministic?

To establish whether the occurrence of interactions is fundamentally stochastic or deterministic, we make a simple prediction: that frequencies of interactions between co-occurring species pairs will consistently be lower than one. To test this prediction, we calculate confidence intervals for the true probability of each interaction in each dataset. Binomial intervals are based on the empirically-observed interaction frequency *µ*, with variance 1/(1-*µ*). While there is some debate as to the best way to calculate confidence intervals for binomial distributions, here we use the Jeffreys interval; the Bayesian credible interval for *µ* with a non-informative Jeffreys prior. This approach yields similarly-sized intervals as the Wilson interval, but with equal tails. These intervals are calculated using the R (R Core Team, 2024) function ‘binom.bayes’ from the package *binom* (Dorai-Raj, 2022). To ensure central intervals using Jeffrey’s prior, we use the argument ‘type=’central” and the default prior shape arguments. Note that while a very large sample size is needed to reliably detect all interactions in a network (Cirtwill *et al*., 2019), our empirical datasets both include sufficient sampling to ensure that sampling effort does not dominate interaction frequencies (Appendix *S3*).

### Are interaction probabilities constant over time and space?

After demonstrating that interactions between co-occurring species tend to occur with *P* (*L_ijy_*) *<* 1, we next test whether *P* (*L_ijy_*) is constant across a set of empirical networks or whether it varies with time and the environment (i.e. whether *P* (*L_ijy_|X_iy_, X_jy_*) is a sufficient model to recreate observed network structure across space and time). To this end, we compare each empirical network to a set of simulated networks where the species in each simulated network are identical to the set of observed species but interactions are included stochastically with a constant probability. Interactions between pairs of co-occurring species that occurred in the metaweb are randomly included or excluded by sampling from a binomial distribution based on the observed mean and variance in interaction frequency for each interaction across all empirical networks. For some networks, this simulation approach often produces very small networks for which some network properties could not be calculated. We therefore exclude any simulated networks that do not include at least two species in each taxonomic group (plants and insects or plants and fungi) and four interactions (i.e., at least one four-species motif).

We calculate number of links, connectance, nestedness, network motif profiles, and species motif roles as described above and then calculate *Z*-scores of the difference between each empirical network and the corresponding set of simulated networks. For number of links, connectance, and NODF these *Z*-scores are straightforward comparisons of the empirical value to the mean and standard deviation of simulated values. For motif profiles and species roles, which are multi-dimensional vectors rather than one-dimensional summaries, we first calculate Bray-Curtis dissimilarity between the empirical motif profile or set of species’ roles and each simulated profile or set of roles using the R (R Core Team, 2024) function ‘vegdist’ from the *vegan* package (Oksanen *et al*., 2024). For motif profiles, we then calculate a *Z*-score based on the mean and standard deviation of Bray-Curtis dissimilarities. For species roles, we calculate a *Z*-score based on the mean and standard deviation of mean dissimilarities across species in each network. In both cases we compare the observed dissimilarities against a baseline of zero for networks with identical motif profiles and species roles to the empirical networks. An absolute *Z*-score greater than 1.96 indicates a significant difference between the empirical and simulated networks. We interpret such differences as evidence that assuming a constant probability for each interaction does not accurately reproduce the empirical network at a given time or place. Note that, because the number of small simulated networks varied substantially between empirical networks, the sample size used to calculate each *Z*-score likewise differs.

### Does accounting for stochasticity in, and environmental imprints on, species and link occurrence improve predictions?

To evaluate how including stochasticity in species and interaction occurrence improves our predictions of observed network properties (number of links, connectance, NODF, network motif profiles, and species motif roles as above), we use scenarios 1 & 2 in Table 1 to simulate 1000 randomisations of each empirical network in both datasets.

**Table 1:**
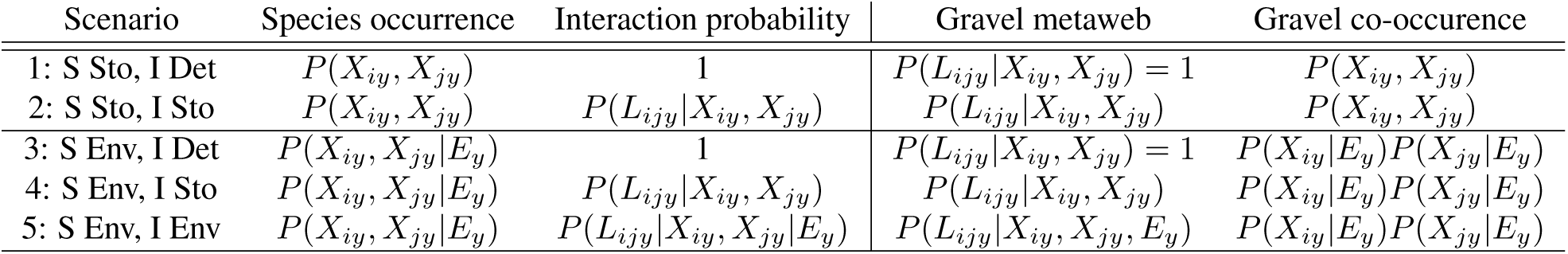
Summary of the scenarios used to simulate ecological networks based on Eltonian and Grinnellian niches, with species (S) and interactions (I) treated either deterministically (Det), stochastically (Sto), or environmentally-dependent (Env). These scenarios partially overlap with those used in Gravel *et al*. (2019); where this is the case, we present the corresponding notation for the metaweb and species co-occurrence from that paper. In all cases, *P* (*X_iy_*) is the probability that species *i* occurs in network *y*, *P* (*L_ijy_*) is the probability that a link occurs between species *i* and *j* in network *y*, and *E_y_* are the environmental conditions of network *y*. In all simulations, only species which were observed to interact in the metaweb and co-occurred in a simulated network were allowed to interact in simulated networks. Note that scenario A was only used when testing whether interaction probabilities are constant over space and time (results shown in Fig. 5).

As above, we discard simulated networks that do not include at least two species in each taxonomic group (plants and insects or plants and fungi) and four interactions (i.e., at least one four-species motif). In all cases, when simulating plant-fungus networks we include the same set of plants as was originally sampled at the focal plot. This avoids confounding variation in network structure due to focal plant selection with variation due to species and link occurrence. We do not create randomisations of the annual pollination networks, as it is unclear how temperature might relate to probabilities of species and interaction occurrence at this scale (e.g., mean annual temperature vs. spring temperature vs. peak temperature).

In scenario 1, species (fungi only for plant-fungus networks; plants and insects for pollination networks) are included or excluded based on random draws from binomial distributions with means equal to the observed frequency of species occurrence across the set of empirical networks. Interactions in the metaweb are assumed to occur deterministically whenever the two species involved co-occurred. In scenario 2, species are drawn as in scenario 1, whereas interactions between locally co-occurring species are assumed to be stochastic. Thus, interactions occurring in the metaweb are included or excluded based on random draws from binomial distributions with means equal to the observed interaction frequency across the set of empirical networks.

Scenarios 3-5 include environmental effects on probabilities of species and/or link occurrence. To evaluate how the probability of occurrence of individual species and individual links varied with the environment, we fit a series of species and link-specific generalised linear models relating occurrence to temperature. For the plant-fungus networks, we use mean annual 2019 temperature extracted from (Parisy *et al*., 2024a,b). For the week-specific observations of plant, pollinator, and plant–pollinator interactions, we use the mean weekly temperature calculated from the hourly measurement of air temperature data extracted from the open data source of Greenland Ecosystem Monitoring (i.e., ClimateBasis; https://data.g-e-m.dk/). Using these temperatures and the set of presences and absences for each species (fungi only for plant-fungus networks) and interactions, we fit binomial models relating occurrence to temperature using the R (R Core Team, 2024) function ‘glm’ from the core package *stats*. When modelling plant-fungus links, we explicitly avoid inflating the number of absences with false negatives due to biased sampling by excluding any observation of the absence of a link if the plant species involved was not sampled at the relevant plot. We model each species and link independently (rather than by a joint model) to avoid circular reasoning (e.g., estimating biotic interactions through residual covariance, then treating the same quantities as responses).

After obtaining models for the probability of each species and link occurring as a function of temperature, we use these models to simulate networks according to scenarios 3-5 in Table 1. In each case, we simulate 100 networks per empirical network for each 0.1^◦^C increment in the observed temperature range across the dataset. In scenario 3, we include or exclude species based on draws from a binomial distribution where the mean is determined by the species-specific temperature function. Metaweb interactions between co-occurring species are assumed to occur deterministically. In scenario 4, species are included or excluded as above and metaweb interactions included stochastically as in scenario 2. In scenario 5, species are included or excluded as above and metaweb interactions between co-occurring species are included or excluded based on draws from binomial distributions where the means are determined by the interaction-specific temperature function.

After simulating randomisations of each network according to each scenario, we then calculate network properties as above. The predictive performance of each scenario is evaluated through visual comparisons with empirical network structure and through statistical tests. For number of links, connectance, and nestedness, these tests take the form of linear regressions predicting the observed values using simulated values, fit using the R (R Core Team, 2024) base function ‘lm’. For motif profiles and species roles, we simply rank the mean Bray-Curtis dissimilarities.

## Results

### Network properties depend on our assumptions regarding Grinnellian and Eltonian niches

When sequentially introducing stochasticity and environmental dependencies to species and link occurrence (Table 1), we observe fundamental changes in network properties. When adding stochasticity to species and interaction occurrence, the connectance of local networks decreased dramatically compared to that of the metaweb, as species and their interactions did not occur in all local network realisations (Fig. 2). Incorporating effects of temperature on species occurrence and interaction probabilities lead to peaks in connectance at certain temperatures; note that the temperatures at which these peaks occurred were not the same when temperature effects on interaction probabilities were included or excluded (bottom panels in columns C and E, Fig. 2). Critically, these toy simulations demonstrate how our predictions about network structural properties depend on the set of observed species and interactions (constant across simulations) *and* the assumptions we make about species and interaction occurrence in local networks.

### Empirical interactions are stochastic, not deterministic

Turning to empirical data, we find strong evidence that across both time and space, interactions are mainly stochastic rather than deterministic (Fig. 4). In the pollination networks, 95% confidence intervals for the true probability of an interaction occurring if both species involved co-occur were generally wide and the mean probability of occurrence was low. Notably, the mean probability of occurrence was lower at shorter time scales (35.8% in weekly networks versus 52.0% in yearly networks), as was the percentage of interactions which occurred in every network where the two species involved co-occurred (7.76% in weekly networks versus 20.5% in yearly networks).

**Figure 4:**
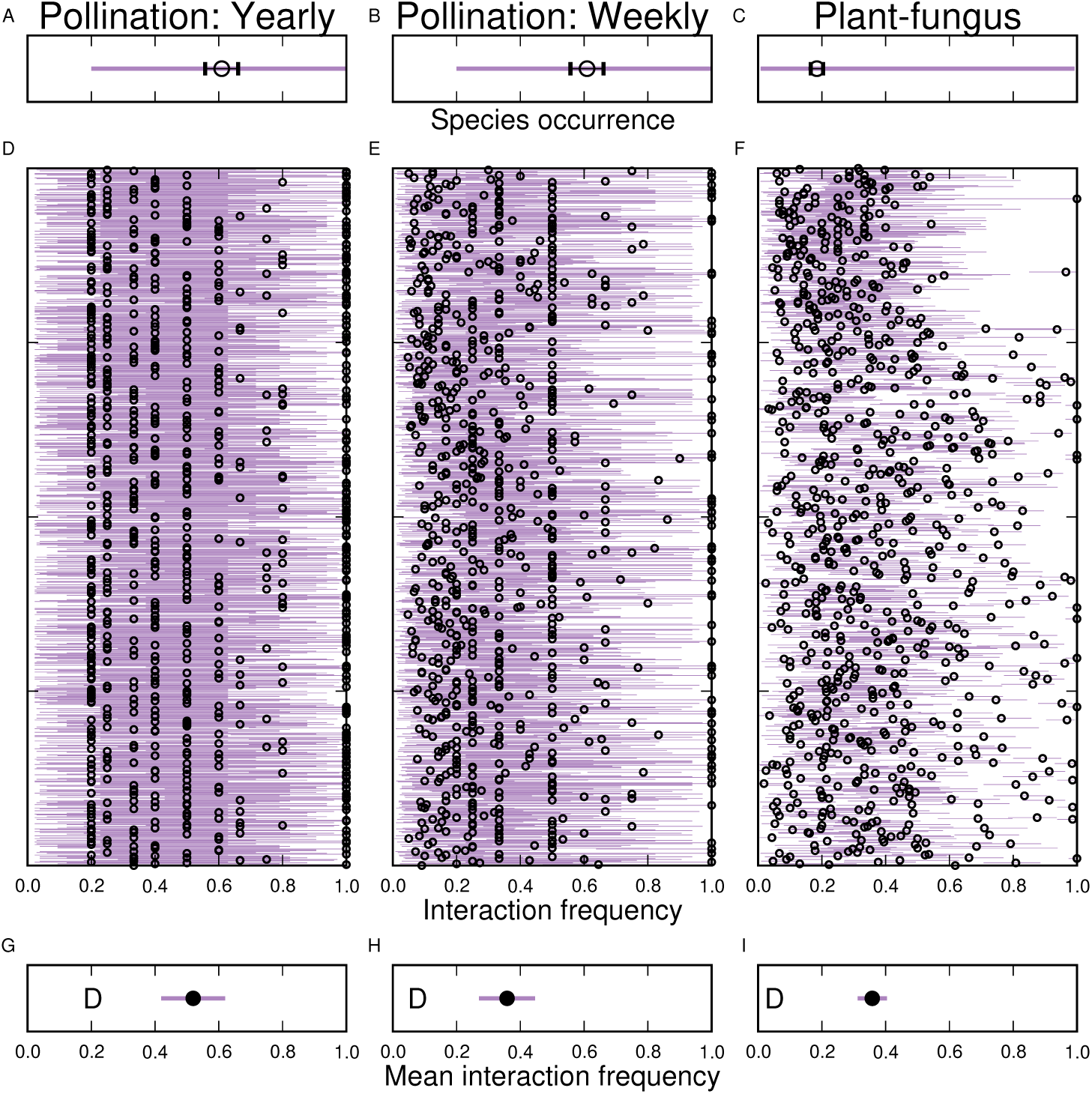
Across both time (left and middle columns) and space (right column), species’ probabilities of occurrence and interaction frequencies are not deterministic in empirical networks. Top: Observed range of interaction frequencies (purple line) with the mean *±*95% CI (dot). Middle: Observed interaction frequencies (circles) and 95% confidence intervals for the true probabililty that an interaction occurs, given the two species involved co-occur, for all interactions in a set of networks. Bottom: Mean interaction frequency (circles) and 5x 95% confidence interval (for visibility) of this mean. The ‘D’ indicates the percentage of apparently deterministic interactions in each network (i.e., those always occurred in observed networks where both species co-occurred). Note that this percentage is much lower at short temporal scales (weekly networks) than long (yearly networks) and is also low at small spatial scales (plots).

In the plant-fungus networks, 95% confidence intervals for the true probability of interaction were generally smaller because of the greater number of empirical networks in this dataset. The mean interaction frequency was similar to the mean frequency in the weekly pollination networks, but the proportion of interactions which always occurred when the member species co-occurred was lower (mean=35.8%, percent deterministic=5.13%). This suggests that there may be substantial variation in fungal communities between individual plants within a species.

### Interaction probabilities vary over space and time

In general, assuming constant frequencies of species and interaction occurrence over space and time (i.,e scenario 1) did not accurately reproduce empirical network structures, even when the species composition of the networks was preserved. While the majority of annual and weekly pollination networks showed statistically similar numbers of links, connectance, and nestedness to the empirical networks, almost all simulated networks had significantly different motif profiles and species motif roles (Fig. 5A, B, D, E, G, H, J, K, M, N). Thus, assuming a constant probability for each interaction does not reproduce the meso-scale structure of a network, even if macro-scale properties can be relatively well preserved.

**Figure 5:**
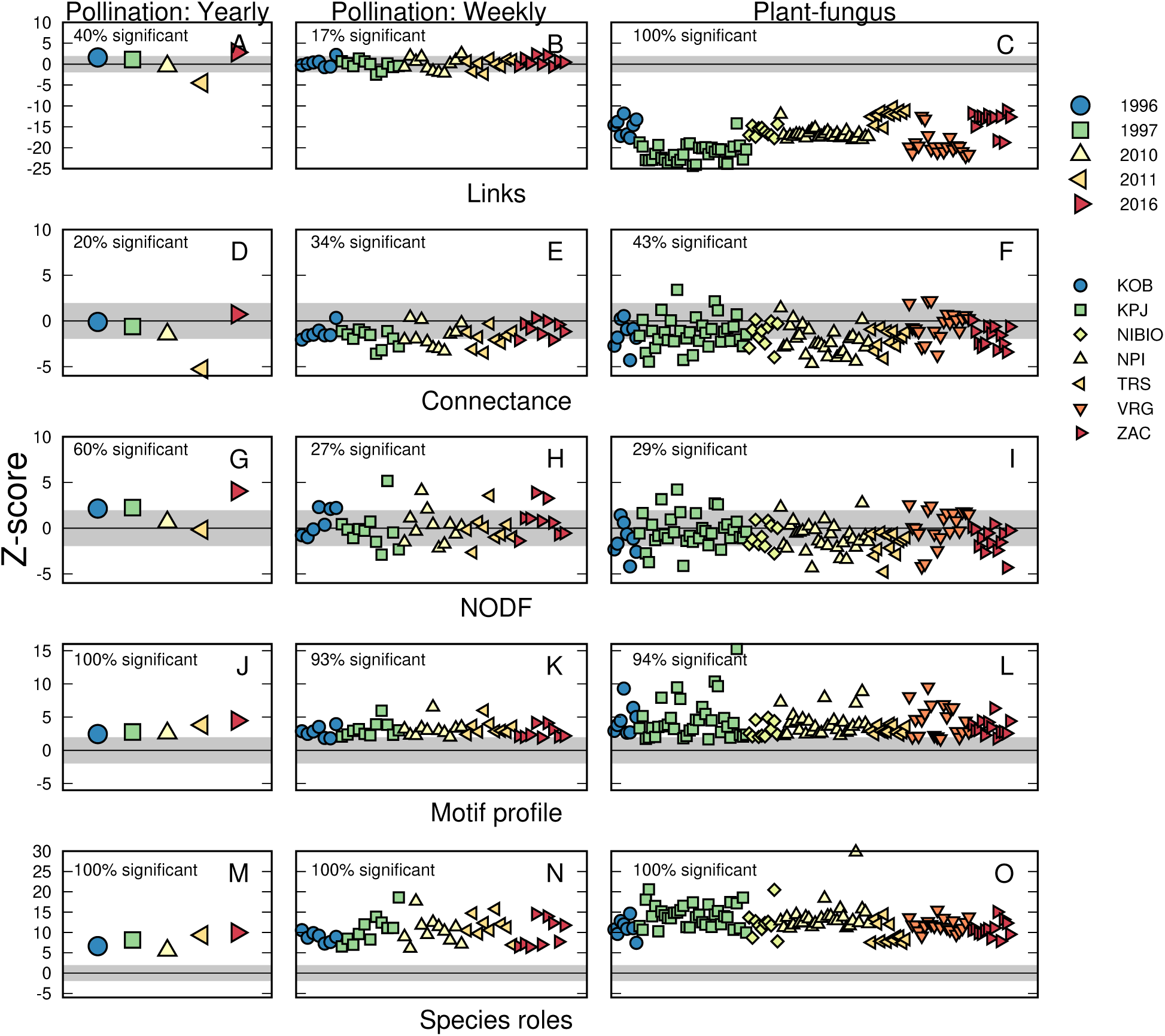
Assuming a constant probability for each interaction can lead to unrealistic simulated networks even when the species composition of networks is preserved. Here we show *Z*-scores of differences between empirical networks and simulated networks including the same species, with each interaction included or excluded based on a binomial distribution using the mean frequency of the focal interaction’s occurrence across all empirical networks in a set. Note that some very small empirical networks produced few simulated networks for which network properties could be calculated; no *Z*-scores are shown for these networks. The proportion of networks shown in each panel with significant *Z*-scores (*p*=0.05) is shown at the top left. For pollination networks, symbol shape and colour indicate year. For plant-fungus networks, symbol shape and colour indicate sampling site, with abbreviations explained in Supplementary Fig.S2.

All of the simulated plant-fungus networks had significantly fewer links than the empirical networks (Fig. 5C). This discrepancy likely occurred because in this system, we have independent data on species co-occurrences and interactions – whereas for the plant–pollinator webs, species are only observed when an interaction occurs (i.e., when an insect sits on a flower). This means that for plant-fungus networks, we have more records of co-occurrences without interactions, and thus a more accurate estimate of true interaction probabilities. At the same time, any simulated network is less likely to include any given interaction without accounting for local factors that make some interactions more likely in particular sites. As some species in the empirical networks had no simulated interactions and were removed from the simulated networks, connectance and nestedness were often preserved (Fig. 5F, I). Network motif profiles and species’ motif roles, however, were almost always different between the observed and simulated networks (Fig. 5L, O).

### Accounting for stochasticity in and environmental imprints on species and link occurrence improves predictions of observed data

Overall, individual species and individual links showed highly variable responses to site-level mean annual temperatures (Fig. 6). In terms of temperature-related patterns in network properties, simulated networks for individual sites (Supplementary Figs. S4-S15, Appendix *S4*) showed fewer links, lower connectance, and often lower nestedness when interactions were treated as stochastic (Scenario X; Table 1) or environmentally-dependent (Scenario X; Table 1) rather than deterministic (Scenario X; Table 1). Treating species and/or interaction probabilities as environmentally-dependent (Scenario X; Table 1) introduced substantial noise into predictions of network properties, but temperature trends were clearest when including temperature effects on species occurrence but treating interactions deterministically (Scenario X; Table 1). This indicates that, in our datasets, effects of temperature on species occurrence and interaction probability often counteract each other.

**Figure 6:**
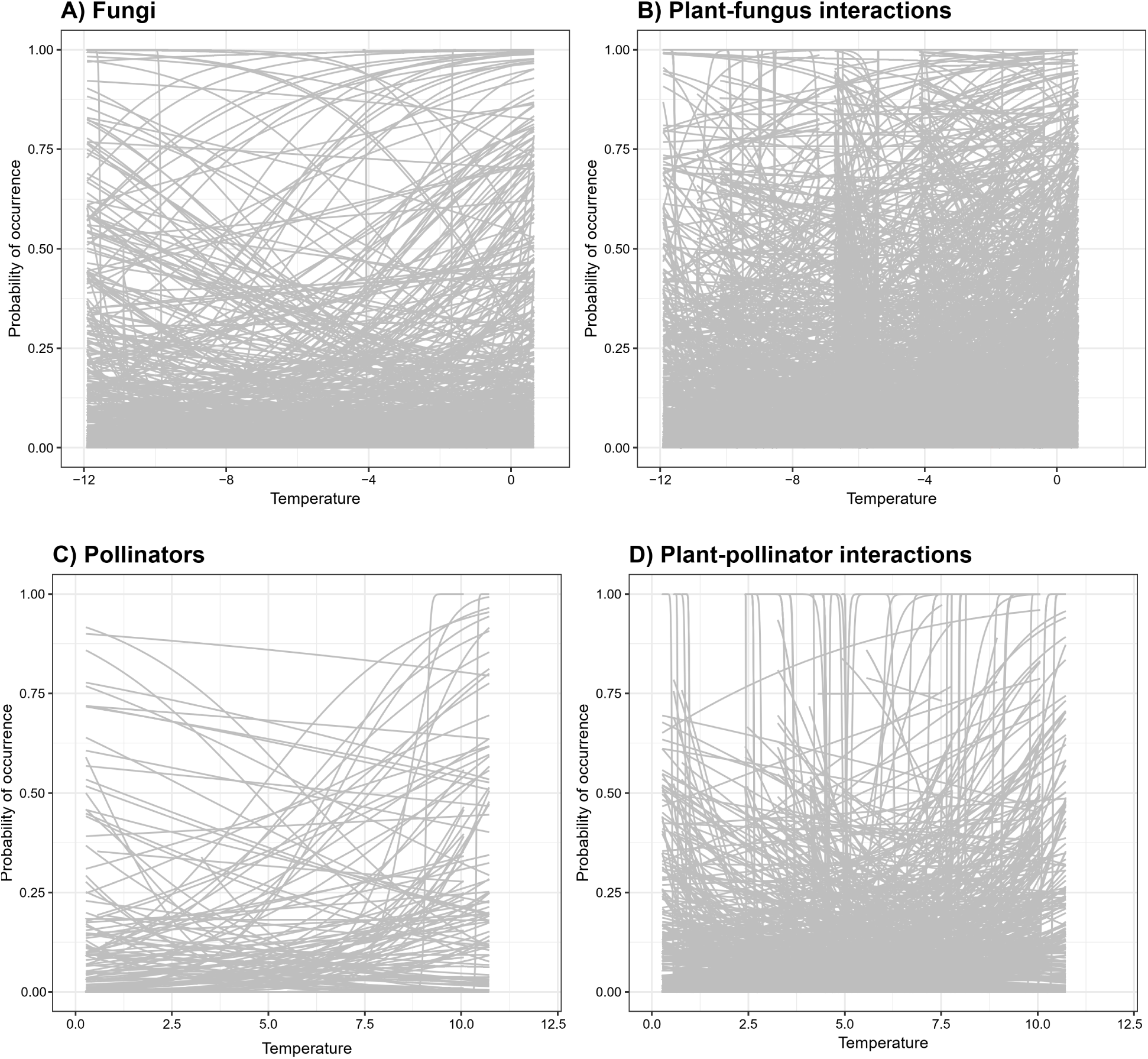
Relationships between interaction probability and temperature are highly variable. We show the individual fitted binomial regression (GLMs) on A) occurrences of individual fungal species, B) individual plant-fungus links, C) occurrences of pollinators species and D) individual pollination links.

In terms of predictive power, adding the assumption that species occurrence is stochastic rather than deterministic substantially improved the fit between simulated and observed network properties (compare scenarios 1 vs. 2 or 3 vs. 4-5 in Table 2), though the pollination networks were poorly predicted by all simulation approaches. Including environmental effects improved predictions of motifs and roles for both datasets and yielded the best (though still poor) predictions for the pollination networks. Although there was substantial variation in the macro-scale properties of simulated networks, the range of simulated properties tended to converge around the observed values when treating interactions stochastically and/or incorporating temperature effects (Figs. 7, 8; S16-S17). For motifs and roles, the variability in dissimilarity between observed and simulated networks increased when interactions were treated stochastically or temperature effects included. Nonetheless, as these dissimilarities tended to be close to 1 when treating interactions deterministically, this increased variability actually implied that at least *some* simulated networks had meso-scale structures similar to the empirical networks.

**Figure 7:**
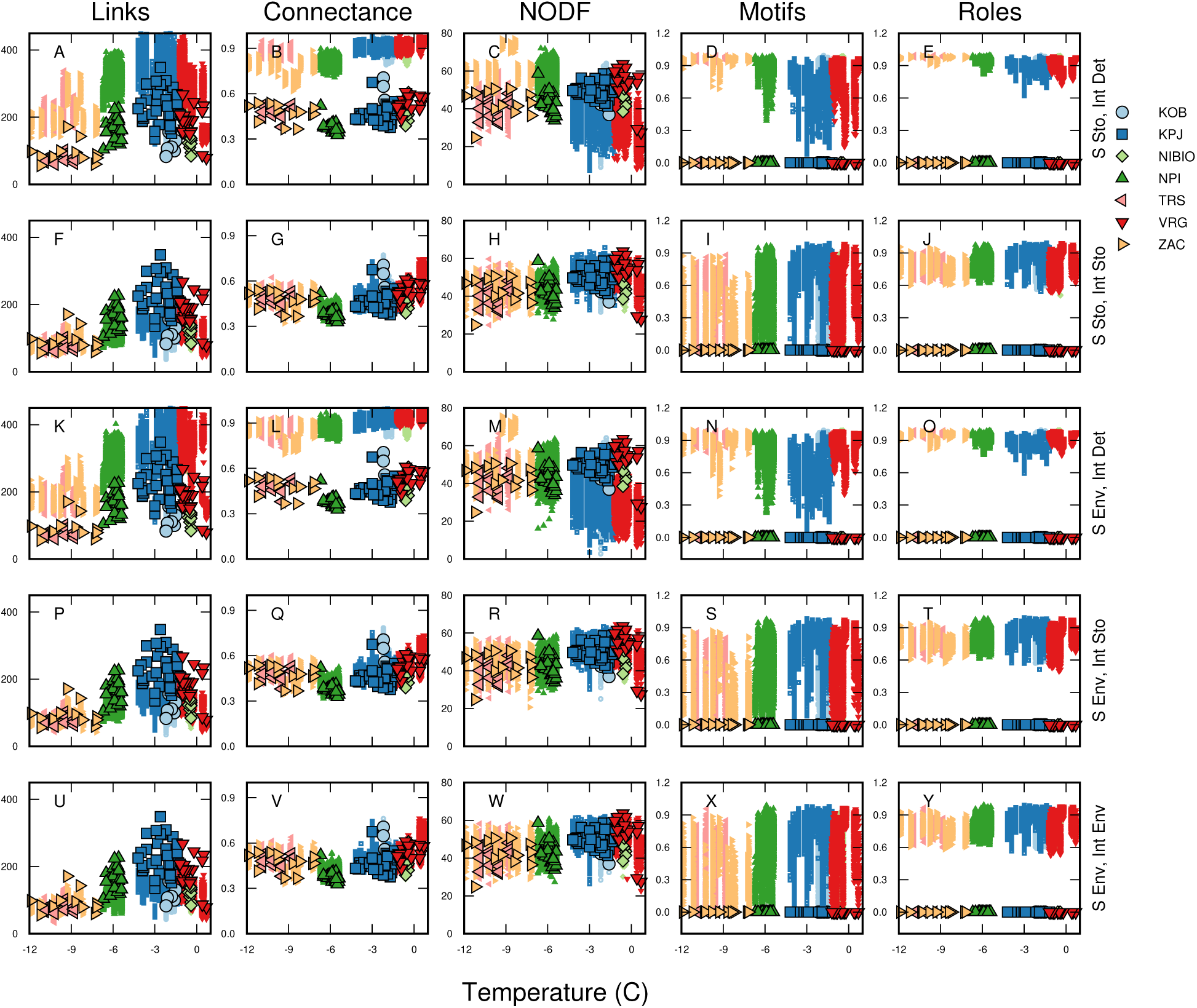
In the plant-fungus networks, including interaction stochasticity leads to simulated networks (small symbols without outline) with summary properties (Links, Connectance, and NODF) more similar to the observed networks (large symbols, outlined in black). For network motif profiles and species roles, which are multi-dimensional measures of network structure, we show the Bray-Curtis dissimilarity between observed and simulated motif profiles or mean Bray-Curtis dissimilarity between observed and simulated species roles. If the observed and simulated networks are identical, this value will be 0 (indicated by large symbols, outlined in black). Although network motif profiles and species’ roles were often quite different between observed and simulated networks, including environmental effects lead at least *some* simulations to approach the observed fine-scale structure of the network.

**Table 2:** Ability of each simulation scenario to capture variation in the observed local networks. For macro-scale properties (links, connectance [C], and nestedness [NODF]), we present *R*^2^ values for linear regressions of observed against predicted values. PF and PP stands for the plant-fungus interaction and plant pollinator data respectively. For multi-dimensional meso-scale properties (motif profiles and species roles), we present estimated mean Bray-Curtis dissimilarities between observed and simulated profiles (Motifs Diff) or roles (Roles Diff). Note that, for motifs and roles, *smaller* differences indicate a better fit.

| Data | Scenario | Links<br>$R^2$ | C<br>$R^2$ | NODF<br>$R^2$ | Motifs<br>Diff | Roles<br>Diff |
| --- | --- | --- | --- | --- | --- | --- |
| PF | 1: S Sto, I Det | 0.538 | 0.242 | 0.165 | 0.838 | 0.937 |
| PF | 2: S Sto, I Sto | **0.604** | **0.645** | **0.272** | 0.548 | 0.804 |
| PF | 3: S Env, I Det | 0.571 | 0.180 | 0.200 | 0.813 | 0.926 |
| PF | 4: S Env, I Sto | 0.603 | 0.635 | 0.260 | **0.545** | **0.802** |
| PF | 5: S Env, I Env | **0.604** | 0.633 | 0.259 | 0.558 | 0.807 |
| PP | 1: S Sto, I Det | 0.001 | <0.001 | <0.001 | 0.873 | 0.970 |
| PP | 2: S Sto, I Sto | <0.001 | <0.001 | <0.001 | 0.723 | 0.916 |
| PP | 3: S Env, I Det | 0.002 | **0.005** | **0.010** | 0.885 | 0.971 |
| PP | 4: S Env, I Sto | <0.001 | **0.005** | 0.006 | **0.701** | **0.899** |
| PP | 5: S Env, I Env | **0.005** | 0.003 | **0.010** | 0.806 | 0.935 |

## Discussion

Understanding the mechanisms behind the assembly and dynamics of ecological interaction networks is key to understanding the biotic consequences of global change (Woodward *et al*., 2010; Tylianakis & Morris, 2017; Pellissier *et al*., 2018). During the last few years, we have studied the extent to which the structure of interaction networks between plants and their partners is driven by environmental conditions. In the current Perspective, we have applied these concepts and methods to the two largest data sets that we have generated from the Arctic. Our analytical framework reveals that both plant–fungus and plant–pollinator networks are stochastic rather than deterministic in nature: of the species and interactions included in the metaweb, only a subset will be observed at a given place and a given time. Further, we find that this probability is not constant over space or time. These effects combine to shape the properties of local network realisations such that networks simulated using deterministic principles are incompatible with empirically observed patterns. However, incorporating temperature effects on occurrence and interaction probabilities still does not universally improve our ability to predict observed network structures, and meso-scale features remain especially hard to capture. Below, we will address the implications of each finding in turn.

### Interactions are stochastic in nature, with probability varying over space and time

In studies of a variety of ecological network types, we have consistently found that species’ probabilities of occurrence and interaction frequencies are not deterministic across either time or space (Gravel *et al*., 2019; Cirtwill *et al*., 2020; Parisy *et al*., 2024a; Fig. 4). Furthermore, assuming a constant probability for each interaction can lead to unrealistic simulated pollination or plant-fungus network properties and meso-scale structures (Table 2), even when the species composition of networks is preserved (Fig. 5). This is especially true for meso-scale structures (motif profiles and species roles). Assuming deterministic interactions leads to mean dissimilarities between observed and simulated structures close to 1 – i.e., nearly as different as possible from the observed. While simulated meso-scale structures were still quite different from the observed when including interaction stochasticity and environmental dependence, there was nevertheless a marked improvement over the deterministic cases.

Acknowledging the stochasticity of interactions between species in future studies is crucially important for three reasons: first, it will dictate what data we can use; second, it will affect how we analyse them; and third, it will determine how we implement these data into predictions of interaction networks under future environmental changes. If we assume that interactions are deterministic, a wide variety of data sources can be used to build a metaweb since “species which have once interacted will always interact” (Lopez *et al*., 2017; Gravel *et al*., 2019). Literature records, expert opinion or diet analyses are then all valid data (Roslin & Majaneva, 2016), regardless of when, where and how frequently an interaction has been observed (cf. Goldwasser & Roughgarden, 1993; Yodzis, 1998; Schlegl *et al*., 2014; Lanuza *et al*., 2025). However, if interactions are stochastic, then the data demands are entirely different. Manifold more data are required to estimate interaction probabilities, since a single observed or inferred interaction provides no information about whether that interaction is common or rare. Even in our two well-sampled datasets, limited observations of each co-occurrence led to wide confidence intervals about the estimates for true interaction probabilities (Fig. 4). A further difficulty is the possibility that interaction frequencies are likely to change over space and/or time depending on environmental factors (Zoller *et al*., 2023). To accurately measure such potential variation, we need many observations of each pair of co-occurring species at each site and time of interest. Simultaneously, data on interaction frequencies should be restricted to the context of the study for which they were collected *or* the data should be analysed with relevant environmental conditions as covariates, as otherwise these data may not be transferable to other locations or times.

Turning from data requirements to analysis, stochastic interactions also imply different connections between the metaweb and local network realisations. If interactions are assumed to be deterministic, then combining records of the species present at a particular place or time with the metaweb will fully determine local networks (Gravel *et al*., 2013; Albouy *et al*., 2019; Cirtwill & Stouffer, 2016). Since interactions are generally stochastic (Fig. 4), this approach generates networks of interactions that *could occur* but do not necessarily *truly occur*. Local network realisations will be chance outcomes of the species present and fundamental interaction probabilities – generating much more variation in local interaction structure (Cirtwill *et al*., 2019). Given this inherent variability, probabilistic measures of network structure should be used in order to distinguish between changes due to normal variation and those that reflect larger underlying trends (Poisot *et al*., 2016).

Predicting how ecological networks may change over time is even more complicated given interaction stochasticity. If interactions were deterministic, a set of species distribution models and the metaweb would be sufficient to predict local networks, assuming all potentially co-occurring species co-occur at least occasionally in the data used to build the metaweb. However, our results show that such an approach does a poor job of reproducing existing network structures (Figs. 7, 8). It is therefore unlikely to accurately predict future networks. Including interactions stochastically provides a closer match to observed network structures, although our results suggest that models assuming constant interaction probabilities are still too simplistic to reliably reproduce network structures (Fig. 5). To understand how network structure is likely to change over space and time requires extensive sampling under a range of environmental conditions. Given the number of samples needed to confidently predict a single interaction probability (Cirtwill *et al*., 2019), this is a tall order. Yet, without such extensive data, environmental dependencies are likely deflated by sampling noise in most empirical data sets – including the current ones (Llopis-Belenguer *et al*., 2023).

### Incorporating temperature effects improves predictions of some – but not all – network properties

For plant–fungus networks, incorporating interaction stochasticicty into network stimulations greatly improved our ability to predict observed network structure (compare scenario 2 to 1 and 4 to 3 in Table 2, Fig. 7). By comparison, the incorporation of explicit temperature-dependencies did not significantly improve predictions of macro-scale network structure. This surprised us, since earlier work has demonstrated strong effects of temperature on the occurrence of plants and fungi (Parisy *et al*., 2024a,b; Barbi *et al*., 2025). By capturing the effect of temperature on species co-occurrence, we had thus expected improved fits. By comparison, predictions of network motif profiles and species roles proved most similar to the observed networks in simulations where species occurrences depended on temperature and interactions were stochastic (Table 2). Still, this gain was mainly relative, since the meso-scale properties of most simulated networks remained quite different from those of the observed networks (Fig. 7).

In the pollination networks, incorporating temperature effects did result in improved predictions of macro-scale *and* meso-scale network properties (Table 2, Fig. 8). Nonetheless, although the distribution of simulated macro-scale properties captured observed properties (Fig. 8), the linear relationship between observed and simulated properties was weak (Table 2). We see two probable reasons for the poor fit in this dataset. First, observed number of links, connectance, and nestedness did not display strong relationships to temperature (*R*^2^=0.036, *R*^2^=0.034, and *R*^2^=0.047 for links, connectance, and NODF, respectively; compare with *R*^2^=0.224, *R*^2^=0.092, and *R*^2^=0.248 for the plant-fungus dataset). This means that there is not much of a relationship between temperature and network structure to capture. Second, for coherence with our analysis of the plant-fungus networks, we modelled temperature as weekly averages and assumed one-directional (binomial) relationships between temperature and probabilities. However, while for the plant–fungus networks temperature effects reflect the impact of site-specific average conditions on the local network, this is not true for the pollination networks. As the latter were sampled over time, temperature effects will combine two imprints: the effect of seasons of different temperatures on plant vs pollinator phenology, and the effect of short-term, contemporary temperature on insect activity during a particular week. From the former perspective, pollination interactions are strongly structured by phenology during the short Arctic summer (Cirtwill *et al*., 2019, 2023). From the latter, temperature does strongly affect the behaviour of high-Arctic plants and invertebrates (Schmidt *et al*., 2023; Tiusanen *et al*., 2019). Our modelling approach will fail to resolve between the two, likely contributing to poor fits. More refined modelling of each component – to be undertaken in the near future – is likely to result in more accurate models. Nonetheless, even our current, simplistic approach to incorporating effects of temperature out-performed models including only stochasticity of species occurrence.

**Figure 8:**
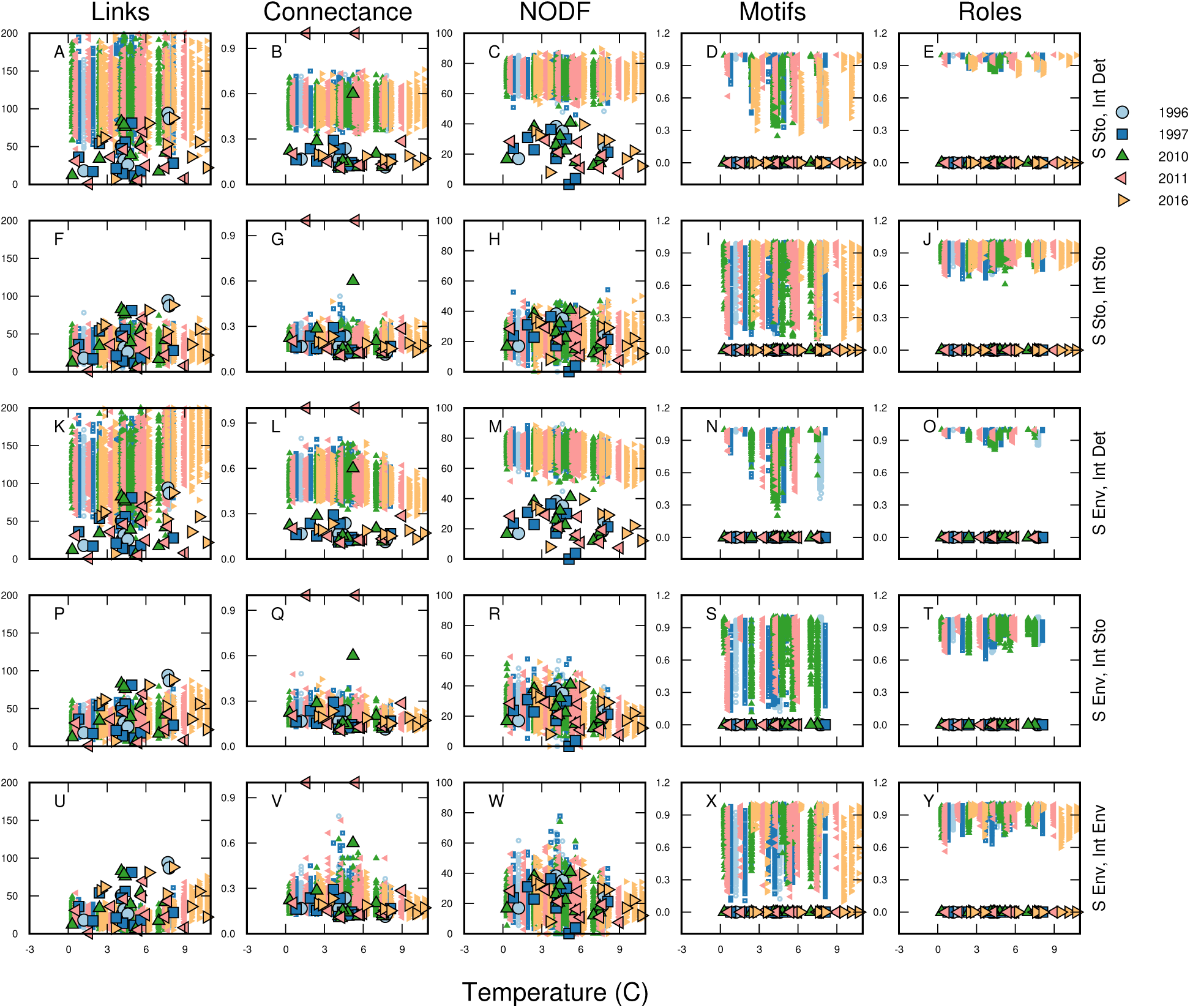
In the weekly pollination networks, including interaction stochasticity leads to simulated networks (small symbols without outline) with more similar summary properties (Links, Connectance, and NODF) to the observed networks (large symbols, outlined in black). For network motif profiles and species roles, which are multi-dimensional measures of network structure, we show the Bray-Curtis dissimilarity between observed and simulated motif profiles or mean Bray-Curtis dissimilarity between observed and simulated species roles. If the observed and simulated networks are identical, this value will be 0 (indicated by large symbols, outlined in black). Although network motif profiles and species’ roles were often quite different between observed and simulated networks, including environmental effects lead some simulations to approach the observed fine-scale structure of the network.

## Conclusions

To arrive at a basic understanding of variation in ecological network properties, and to derive and validate predictions for their future change, we must start from two pieces of information: the fundamental nature of interactions (stochastic or deterministic), and their environmental dependency. In our work, we have found that both species and link occurrence are stochastic, with probabilities conditional on the environment; that these environmental dependencies scale to the network level; and that introducing stochasticity and environmental dependency substantially improves the match between simulated and empirically observed patterns in network properties (as compared to assuming that interactions occur deterministically whenever the species involved co-occur). Altogether, we therefore argue that research based inferring local network structure from a single, deterministic metaweb should be abandoned. Instead, we should at least quantify the probabilities of species and interaction occurrences over space and time. Much preferably, we should quantify environmental impacts on both species and links, thus merging the Grinnellian and the Eltonian niche concept.

## Author contribution

TR conceptualized the study. BP, AC and TR designed the statistical analyses. AC analyzed the data, with input from BP and TR. BP and AC performed the visualisation with input from TR. BP, AC and TR wrote the first draft of the manuscript. All authors then helped revise the manuscript or provided comments on the final manuscript.

## Acknowledgments

TR was funded by the European Research Council (ERC) under the European Union’s Horizon 2020 research and innovation programme (ERC-synergy grant 856506—LIFEPLAN). Data collection for plant-fungus networks was supported by the Academy of Finland (grant 322266 to TR) and an INTERACT Transnational and Remote Access grant (Call 2020, project VEGA). The Greenland Ecosystem Monitoring intiative is thanked for access to temperature data from Zackenberg. We acknowledge CSC-IT Center for Science Ltd., Espoo, Finland, for the allocation of computational resources.

## Authors Biography

Dr. Bastien Parisy is a postdoctoral researcher at the University of Helsinki, Finland. Dr. Alyssa Cirtwill is CEO of the consultancy company Carex Ecologics, Canada, specialized in data analyses. Prof. Tomas Roslin is a professor at the Swedish University of Agricultural Sciences in Uppsala, Sweden, and the University of Helsinki, Finland. All authors are community ecologists studying ecological networks formed by plants and associated organisms.

## Data availability statement

The data of plant—pollinator interactions from Zackenberg, Greenland, were adopted from Olesen *et al*., 2008; Rasmussen *et al*., 2013; Cirtwill *et al*., 2023. The data of plant-fungus networks were adopted from Parisy *et al*., 2024a,b. Codes and data are privately provided for peer review and will be made publicly available upon acceptance of this manuscript.

