## Appendix for "The environmental dependence of ecological interaction networks"

#### Table of contents

- S1: Setting the stage: simulating ecological networks under different scenarios of Eltonian and Grinnellian niches
  - Methods and results used to construct Fig. 2, Main Text (reproduced as Fig. S1)
- S2: Empirical data
  - Details of sampling and map of sites (Fig. S2)
- S3: Does sampling effort dictate interaction frequencies?
  - Methods and results (Fig. S3) testing whether observed interaction frequencies are strongly constrained by the number of observations per interaction
- S4: Simulated properties of empirical networks at different temperatures
  - Figs. S4-S15 showing properties of simulated networks across the observed temperature range
- S5: Alternate display of observed and simulated network properties
  - Figs. S16-S17 showing simulated network properties plotted against observed values

### S1: Setting the stage: simulating ecological networks under different scenarios of Eltonian and Grinnellian niches

#### Methods

To characterise variation in ecological interaction networks under different scenarios of environmental forcing (Fig. S1), we conducted explicit scenario analysis. In brief, we compared the relationship between a single environmental gradient (here: temperature) and expected patterns in and network structure under six scenarios, moving from a fully deterministic model ("all species present everywhere, all links occur when they occur in the metaweb") towards scenarios adding stochasticity to species occurrence and link occurrence, and finally introducing environmental dependence of occurrence probabilities of both species and links (Table 1, Main Text).

To relate this approach to previous contributions, we note that our scenarios are similar to the sequential exploration conducted by Gravel *et al.* (2019). Following their notation, we use $\mathrm{X}_{iy}$ to represent the probability that a species $i$ co-occurs at location $y$ – given that species $i$ is a member of the community (i.e., occurs in the metaweb). For a deterministic representation of the community, $X_{iy}$ equals one across all sites $y$. For a probabilistic representation, we define $X_{iy}$ as a stochastic process taking a value of 1 when the species occurs at site $y$ and a value of 0 when it is absent. In a similar vein, we use $L_{ijy}$ to represent the probability that an interaction between species $i$ and $j$ occurs at location $y$ – given that these species have ever been observed to interact (i.e., they interact in the metaweb). For a deterministic representation of interactions, $L_{ijy}$ equals one whenever species $i$ and $j$ co-occur. For stochastic representations, we define $L_{ijy}$ as a stochastic process taking a value of 1 when an interaction occurs and a value of 0 when it does not.

After simulating networks with deterministic or stochastic species and interaction occurrences, we explore the probability that species $X_i$ and $X_j$ and their interaction $L_{ij}$ all occur in network $y$, conditional on the set of environmental conditions $E_y$: $(P(X_{iy}, X_{jy}, L_{iyj}|E_y)$. As noted by Gravel *et al.* (2019), this probability can be decomposed into two parts using the product rule of probabilities: $P(X_{iy}, X_{jy}, L_{iyj}|E_y) = P(X_{iy}, X_{jy}|E_y)(L_{iyj}|E_y)$ Here, the first term on the right-hand side of the equation is the probability of observing the two species co-occurring at location $y$. It corresponds to the Grinnellian dimension of the niche. The second term represents the probability that an interaction occurs between species $i$ and $j$, given that they are co-occurring. In the scenarios where this probability is conditional on the environment, we introduce environmental dependence ($|E_y$; Table 1 Main Text).

We illustrate these simulations in a toy metaweb in Figure S1 (column A, top). In a fully deterministic scenario ($X_{iy}$ fixed to one), all species occur at all temperatures (middle) and all possible links (filled squares) always occur (probability of occurrence = 1, indicated by black fill). Here, all local realisations of the network are identical to the metaweb and simulations are not necessary. In the first simulation scenario (Fig. S1, column **B**), we introduce stochasticity in species occurrence (making $X_{iy}$ a stochastic event). That is, each species in the network (identified by green squares or purple circles) occurs with a certain probability (indicated by the matching-coloured line in middle) that is constant across temperatures. To maintain comparability with later scenarios, these probabilities are the mean of probabilities of occurrence at each 1°C temperature increment used in later simulations.

In the third scenario (Fig. S1, column **C**), we begin to introduce environmental effects on the network. Each species' probability of occurrence now varies with temperature (with each species having an optimum temperature where its probability of occurrence is 1 and a range of temperatures with lower probabilities of occurrence). The probability of occurrence for each species at each temperature was defined as:

$$p = 1 - [|(optimum - temperature)| * specialsation]^2 \tag{1}$$

Optimum temperatures ranged from 4-26°C and specialisation parameters ranged from 0.05-0.125 (higher values produce narrower temperature ranges). We note that this approach produces species with unimodal, quadratic responses to temperature. Linear or multi-modal responses could also be possible but are left as an exercise for the enthusiastic reader.

In the fourth scenario (Fig. S1, column **D**), we begin to introduce variability in interaction probabilities. Each interaction occurs (assuming the species involved co-occur) with a certain probability that is constant across temperatures. For comparability with other scenarios, and similar to species occurrence probabilities in **B**, these probabilities are the means of temperature-dependent probabilities calculated at 1°C increments in **E**.

Finally, we introduce environmental effects on the probability that each link occurs. As with species, each link was assigned an optimum temperature (ranging 3-28°C) and a level of temperature specialisation (ranging 0.05-0.25). The probability of occurrence for each link was calculated at each 1°C temperature increment following equation 1.

#### Results

Our set of scenarios sequentially introducing stochasticity and environmental dependencies to species and link occurrence (Table 1, Main Text) demonstrates fundamental changes in network properties. In the fully deterministic scenario (Fig. S1A), all species occur at all temperatures (middle panel from top) and all possible links always occur (probability of occurrence = 1, indicated by black fill in top panel). As a result, the connectance of this network is constant across the entire temperature range (bottom panel).

In the second scenario (Fig. S1B), we introduced stochasticity in species occurrence. In this scenario, links always occur if the two species involved co-occur (links remain deterministic), but because species pairs have different probabilities of co-occurrence (varying shades of grey in top panel), links appear with different frequencies in realised occurrences of the network. Thus, varying probability of occurrence between species results in variable connectance (mean and standard deviation shown in bottom panel) which does not vary consistently over temperature. Note that introducing stochasticity in species occurrence leads to substantially lower connectance than in the deterministic network, because many interactions do not occur when the species involved do not co-occur (compare bottom panels in Fig. S1A and B).

In the third scenario (Fig. S1C), we introduced environmental effects on the network. In this scenario, the probability of co-occurrence for each species pair varies with temperature (compare darkness of filled cells in top panel across example temperatures). Although interactions are once again assumed to occur whenever the species involved co-occur, as in Fig. S1A-B, the temperature dependence of species means, in this case, that species are more likely to co-occur and interact at intermediate temperature values. Although the mean connectance across temperatures is now the same as that in Fig. S1B (as the probability of occurrence used for each species is the mean of its probability of occurrence at each 1°C temperature increment), both the mean and variability of connectance vary strongly across temperatures (bottom panel).

In the fourth scenario (Fig. S1D), we introduced stochastic variability in interaction probabilities. Because these probabilities are constant over temperature but lower than 1, the net effect is to dampen the influence of temperature on the probability that a given interaction occurs (compare more-similar shading in cells across example temperatures in the top panel with the dramatic differences in the top panel of Fig. S1C). This results in a more subtle peak in connectance at intermediate temperatures (bottom panel). Because interactions are no longer assumed to always occur if the species involved co-occur, the mean connectance in simulated networks is also lower than in scenario C (compare bottom panels of Fig. S1C-D).

Finally, we introduced environmental effects on the probability that each link occurs. As with species (Fig. S1C), this approach produces unimodal quadratic relationships between probability of occurrence and temperature (Fig. S1E; other types of relationships are possible). In this example, interactions are generally more likely to occur close to the temperature optima of at least one of the species involved. At the network level, this produces probabilities of occurrence that are more similar to those in scenario C than the constant probabilities in D (compare example networks in top panels of Fig. S1A-E). However, because no interactions are especially likely to occur at intermediate temperatures (Fig. S1E, middle panel), the relationship between connectance and temperature is bimodal rather than unimodal (Fig. S1E, bottom panel).

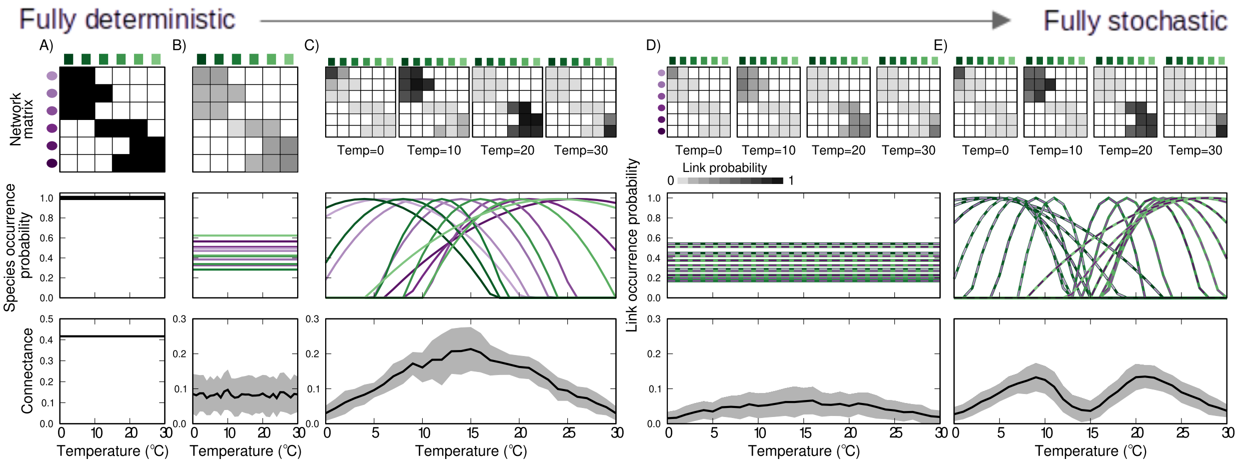


**Figure S1: Reproduction of Fig. 2, Main Text.** The stochastic vs deterministic view on species interactions. Vertical panels show impacts on different aspects of network structure as we horizontally move from a deterministic view to one which incorporates Grinnellian niches of species and their interactions. The top row shows the network matrix, in which cell shading indicates the probability that a link occurs, the middle row shows the probability of each individual species (A-C) or link (D-E) occurring at each temperature, and the bottom row shows the mean (±SD) connectance of 100 simulated observed networks for each temperature. (Note change in scale of the connectance axis between A and B–E.) Impacts are compared among six scenarios: (A) Species occur with probability 1 at all temperatures, and feasible interactions always occur. (B) Species occur with a constant probability, which differs between species. This reduces the probability that interacting species will co-occur and interact, leading to lower, and variable, connectance. (C) Species have unique Grinnellian niches with respect to temperature (curves in middle panel), leading to varying probabilities of co-occurrence. As in B), species always interact when they co-occur with a feasible partner. Columns D–E combine these Grinnellian niches with interaction stochasticity: (D) Interactions are stochastic with constant probability across temperatures, lowering the probability of any interaction occurring and decreasing the connectance of the resulting observed networks. (E) Some interactions are more likely at cooler temperatures, others at warmer temperatures (depending upon the temperature optima of the species involved). In this example, this results in somewhat higher connectance at moderately low (5-10°C) and moderately high (20-25°C) temperatures than when interaction probabilities are constant across temperatures (D). Note that the probabilities of link occurrence in D are the average values of the curves in E; likewise, the probabilities of species occurrence in B are the average values of the curves in C. For simulation methods, see Appendix *S1*.

### S2: Empirical data

As a basis for analyses of empirical data sets, we use the two largest sets of systematically collected data on ecological interaction networks of which we are aware. To resolve variation in time, we use data on plant—pollinator interactions from Zackenberg, Greenland, collected at a weekly resolution during five different years (1996, 1997, 2010, 2011, and 2016; Olesen *et al.*, 2008; Rasmussen *et al.*, 2013; Cirtwill *et al.*, 2023). To resolve variation in space, we use data on plant—fungus interactions collected across the Arctic during the summers of 2020 and 2021 Parisy *et al.* (2024a,b).

As described in Olesen *et al.*, 2008; Rasmussen *et al.*, 2013; Cirtwill *et al.*, 2023, flower-visitor networks in Zackenberg were characterized within a constant 500 × 500-m plot (Fig.S S2). Each year, the flower-visitor networks were characterized by identifying all insects visiting two individuals of each focal plant species occurring within the plot, during a 40-min observation on each day with fine weather during the entire snow-free season. Flower-visitors were identified morphologically to the finest possible taxonomic level, but also collected and further identified using DNA barcoding to confirm taxonomical affiliation. In total, the plant-pollinator networks were aggregated at the level of a) weeks (n=49) and b) years (n=5) (Fig.S S2). For full details, see Olesen *et al.*, 2008; Rasmussen *et al.*, 2013; Cirtwill *et al.*, 2023.

As described in Parisy *et al.*, 2024a,b, a total of 1450 root and 1450 soil samples associated with 12 targeted plants were collected across a gradient of 14.5 latitude during the summers of 2020 and 2021 (Fig.S S2). Within each sites, at least three local transects were defined. These transects were located at least 250 meters apart from each other across a joint elevation gradient. Within three sites (Kilpisjärvi, Varanger, and Zackenberg), the sampling was more intensive and followed a stratified random sampling design across multiple local elevational gradients. Each transect included a minimum of four plots, each with a radius of 25 meters containing five locally prominent plant species (out of the 12 targeted species). The root associated fungi and the fungal soil communities of each plants species were characterized through eDNA metabarcoding (see Parisy *et al.*, 2024a,b for the full details.). In total, the plant-fungus networks were aggregated at the level of a) plots (n=129 and b) sites (n=7). From these empirical networks, we constructed our simulated networks using the same five plants collected within a site. To these plants, we randomly selected the same number of fungi as in the observed network.

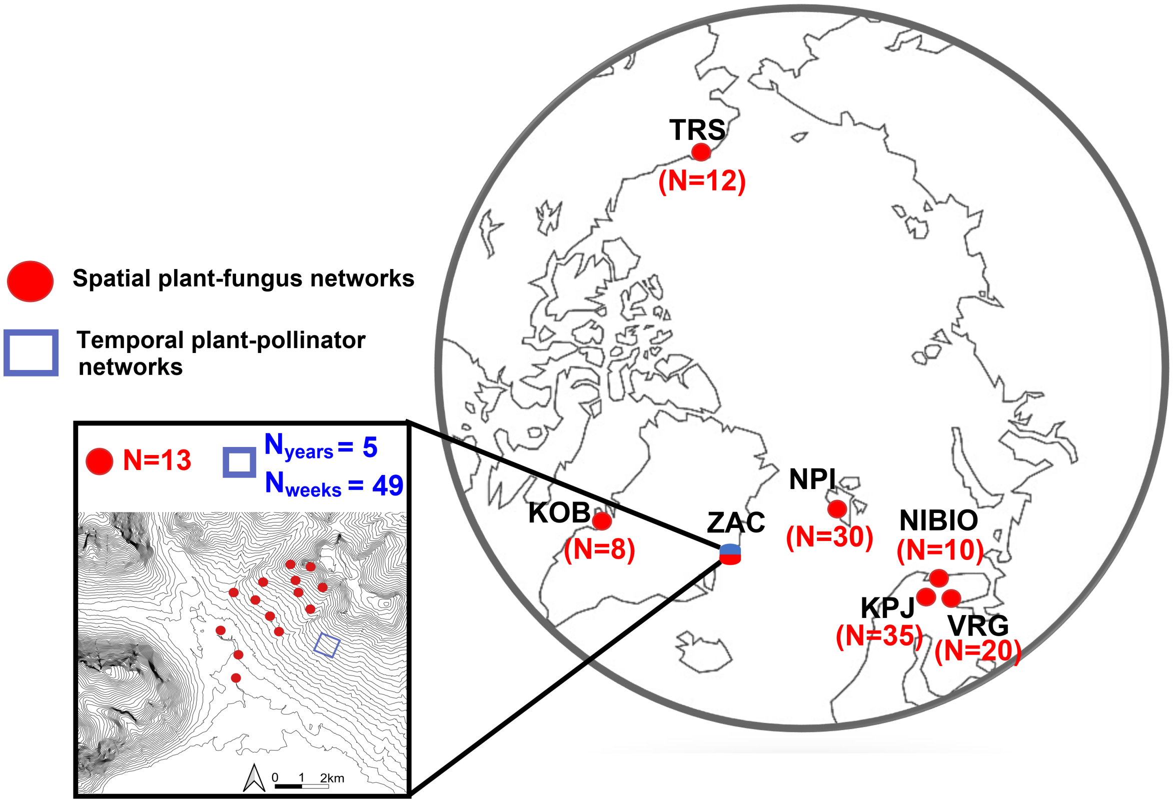


**Figure S2:** Study area. In the global map, each dot colour corresponds to a specific dataset. Plant fungus networks characterized across the Arctic in red and plant-pollinator networks characterized across time. The inset represent the Zackenberg valley where we have characterized both types of networks. (Note that the dots in the inset are not drawn to scale.) The number N corresponds to the number of characterized networks, which refers to, whether the number of plots for the plant fungus networks in red, whether the number of years (i.e, Nyears) and weeks (i.e., Nweeks) of which the plant-pollinators have been characterized. Site-specific acronyms:KPJ= Kilpisjärvi, Finland; VRG= Varanger Peninsula, Norway, NIBIO= Gandvik valley, Norway; KOB= Kobbefjord, SouthWest Greenland; TRS= Toolik Research Station, USA; NPI= Norsk Polarinstitut, Ny Ålesund, Svalbard; ZAC= Zackenberg, North-East Greenland.

### S3: Does sampling effort dictate interaction frequencies?

Our approach to testing whether interactions are deterministic or stochastic relies on an explicit assumption: that no sampling noise is added to the observed probability, i.e., that an interaction will be detected whenever it occurs. This is a strong assumption, since $n$ species can, in principle, interact in $n^2$ different ways. Thus, for a species-rich community, (very) large sample size will be needed to reliably reveal each interaction (Cirtwill *et al.*, 2019). As our samples are naturally finite in nature, we thus added an additional validation by ranking interactions by their observed frequencies. Since more frequent associations should be observed with less noise, we expected the most common interactions to approach a value of one – should interactions be deterministic. Instead, fitting binomial regressions relating interaction probability to the number of observed co-occurrences (with the R (R Core Team, 2024) base function 'glm') revealed very little of a relationship (pseudo-$R^2$ ranging 0.030-0.088).

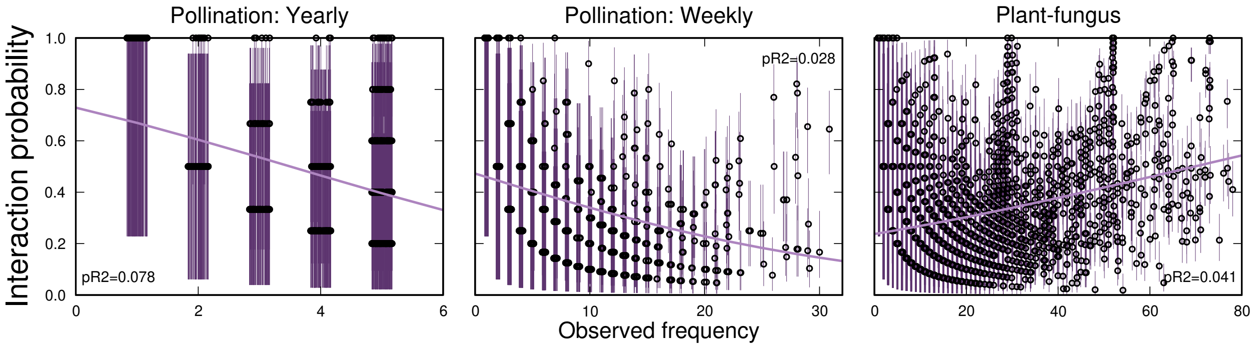


**Figure S3:** If limited sampling constrained interaction frequencies, we would expect that increasing numbers of observed co-occurrences would lead to increasing interaction probabilities, with probabilities approaching 1 for the most frequently-observed co-occurrences. In both of our datasets, we observe a wide range of interaction probabilities even among the species most often observed co-occurring.

### S4: Simulated properties of empirical networks at different temperatures

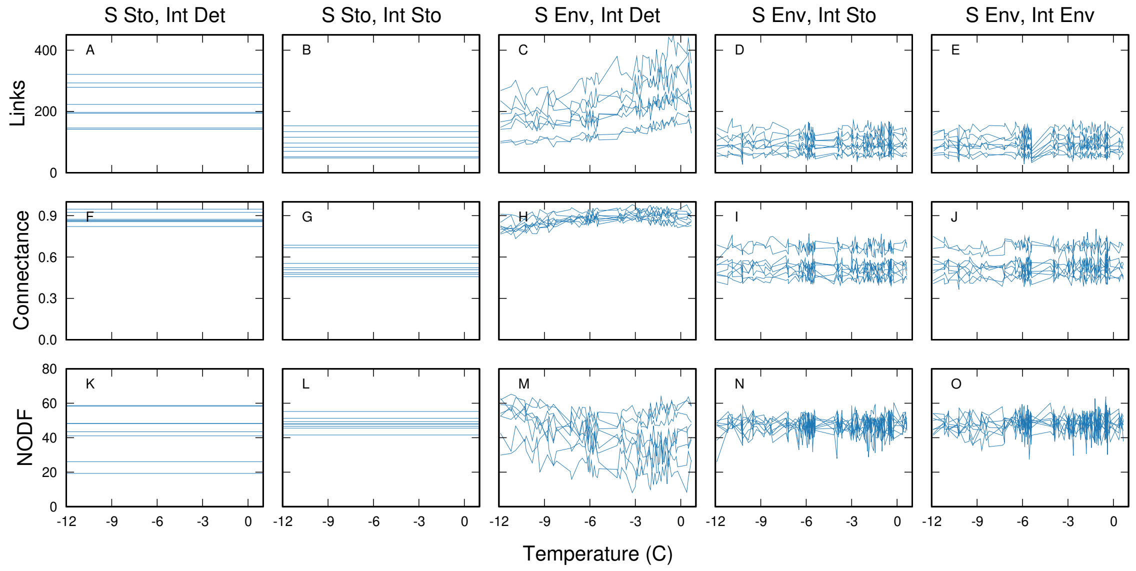


**Figure S4:** Properties of simulated plant-fungus networks over the entire observed temperature range, KOB plots shown. Networks were simulated for each 0.1 degree increment within the range shown.

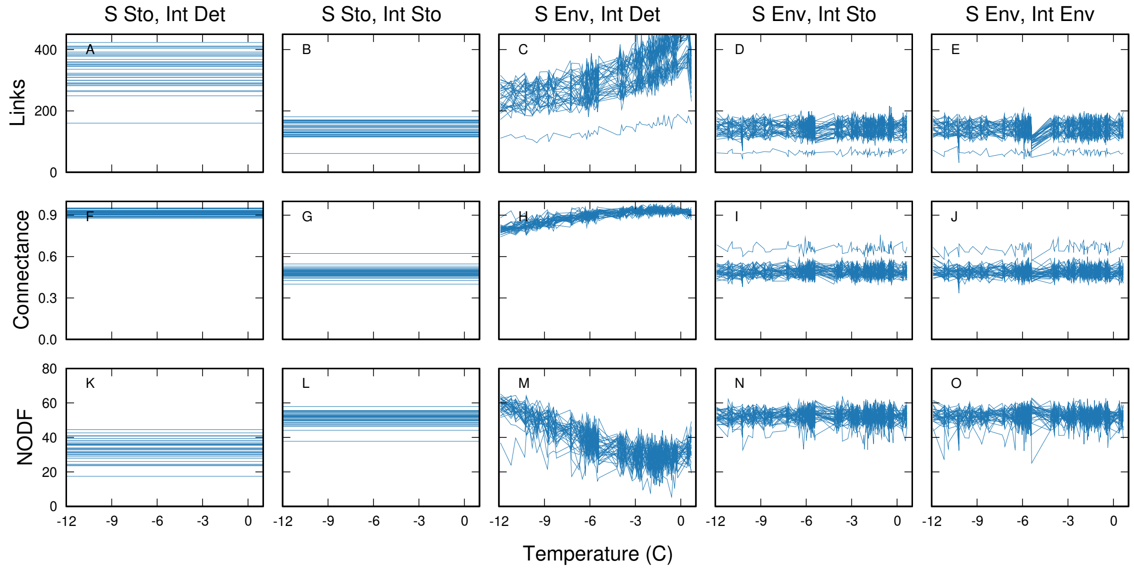


**Figure S5:** Properties of simulated plant-fungus networks over the entire observed temperature range, Kilpisjärvi plots shown. Networks were simulated for each 0.1 degree increment within the range shown.

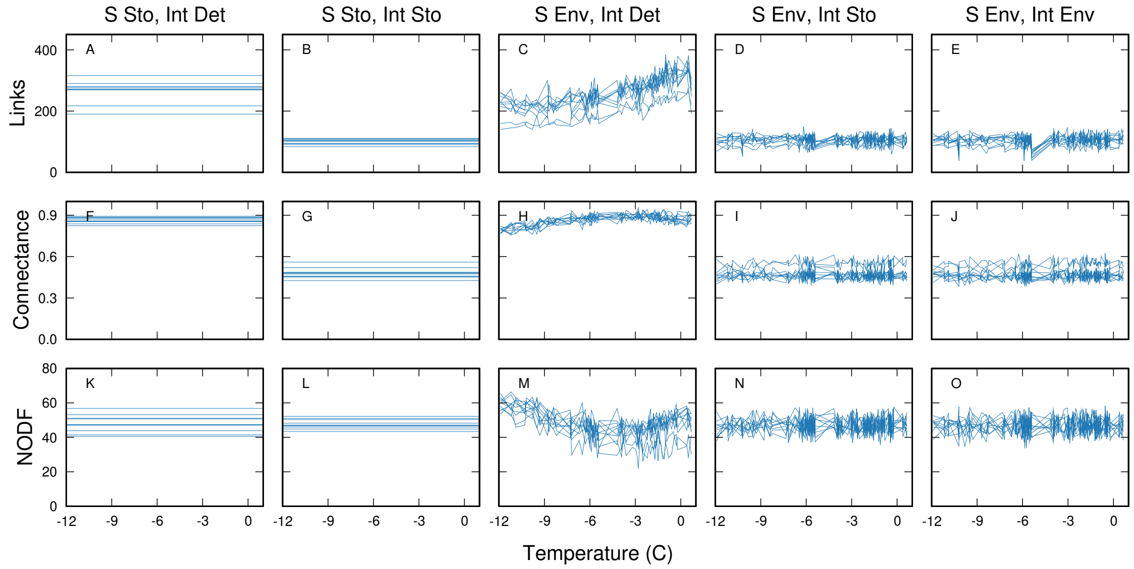


**Figure S6:** Properties of simulated plant-fungus networks over the entire observed temperature range, NIBIO plots shown. Networks were simulated for each 0.1 degree increment within the range shown.

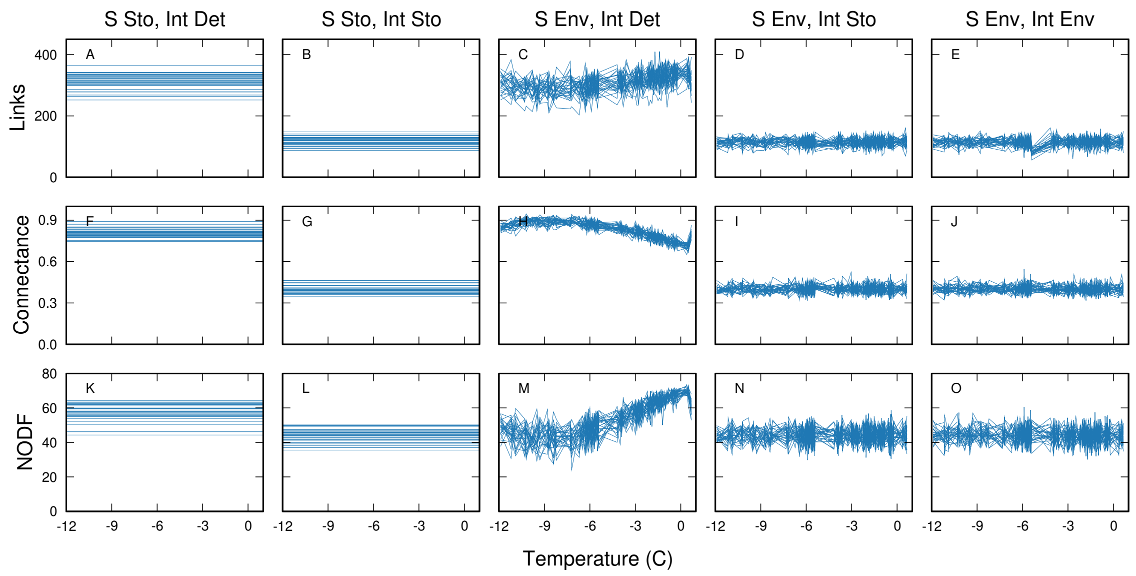


**Figure S7:** Properties of simulated plant-fungus networks over the entire observed temperature range, NPI plots shown. Networks were simulated for each 0.1 degree increment within the range shown.

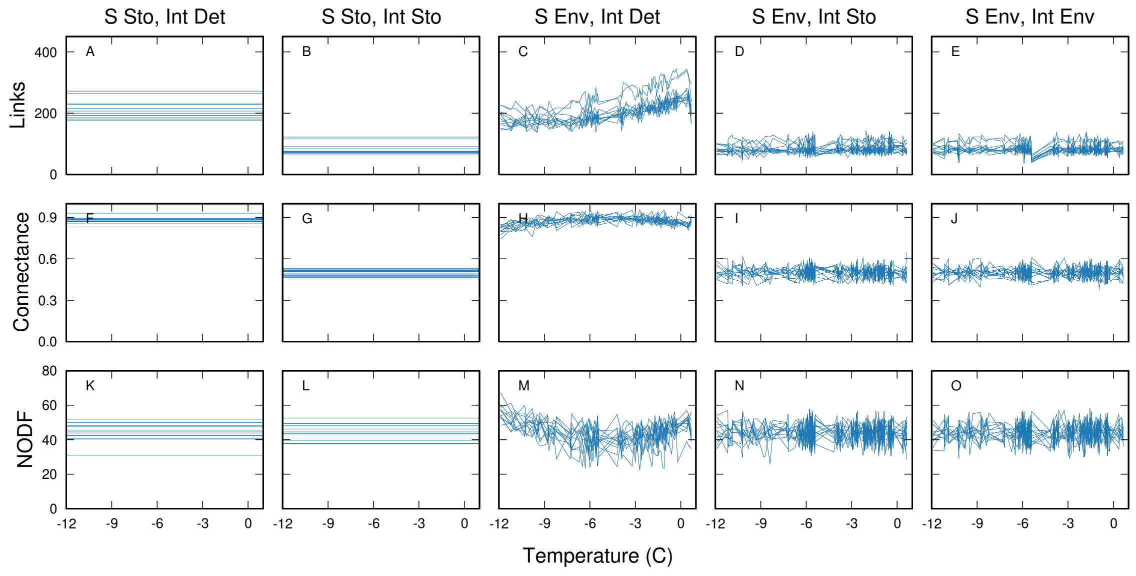


**Figure S8:** Properties of simulated plant-fungus networks over the entire observed temperature range, TRS plots shown. Networks were simulated for each 0.1 degree increment within the range shown.

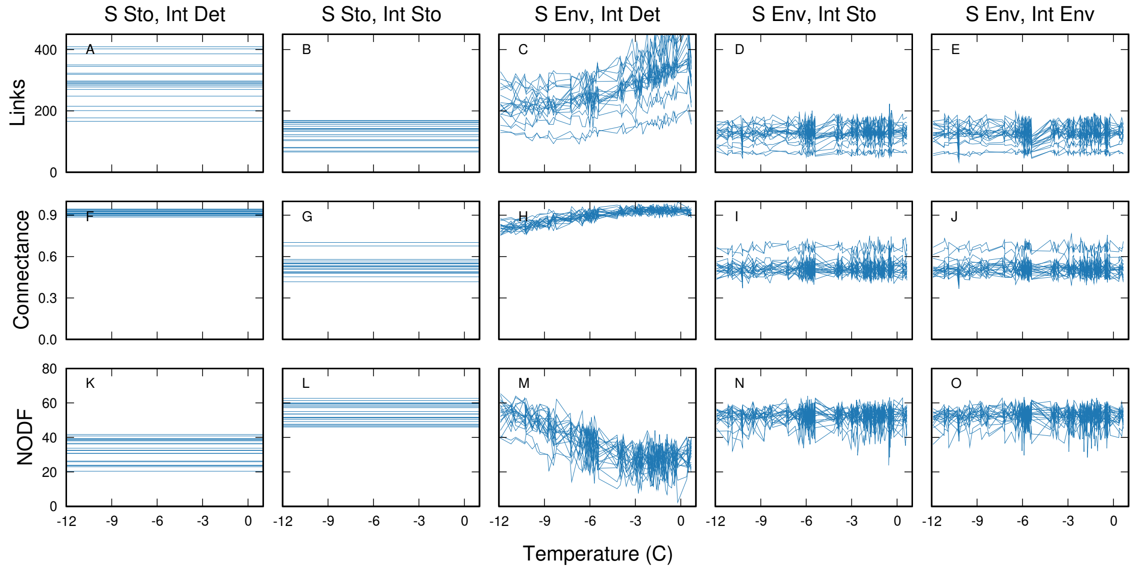


**Figure S9:** Properties of simulated plant-fungus networks over the entire observed temperature range, Varanger plots shown. Networks were simulated for each 0.1 degree increment within the range shown.

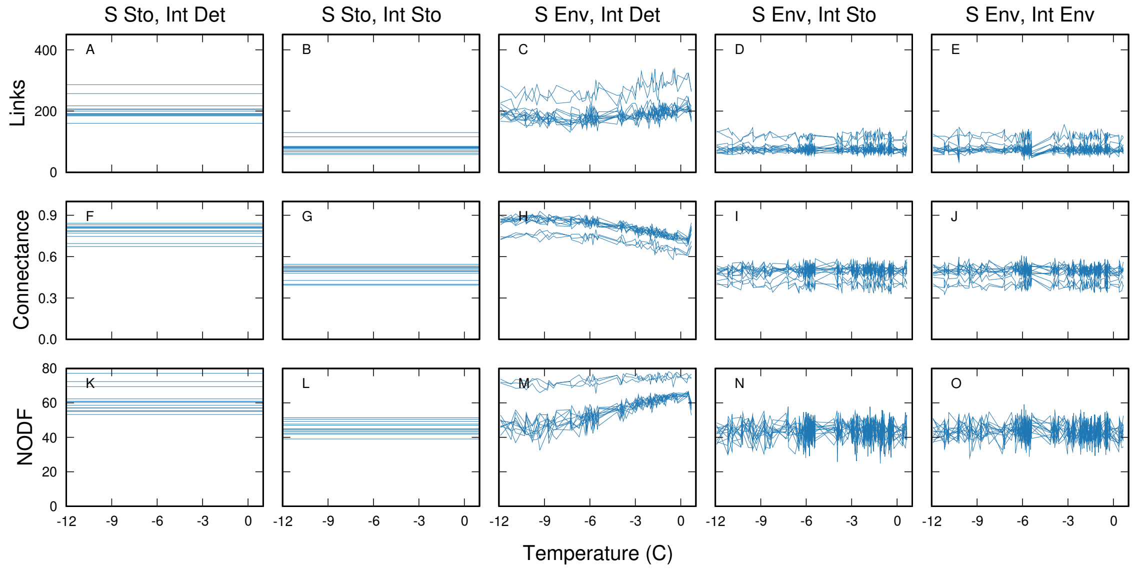


**Figure S10:** Properties of simulated plant-fungus networks over the entire observed temperature range, Zackenberg plots shown. Networks were simulated for each 0.1 degree increment within the range shown.

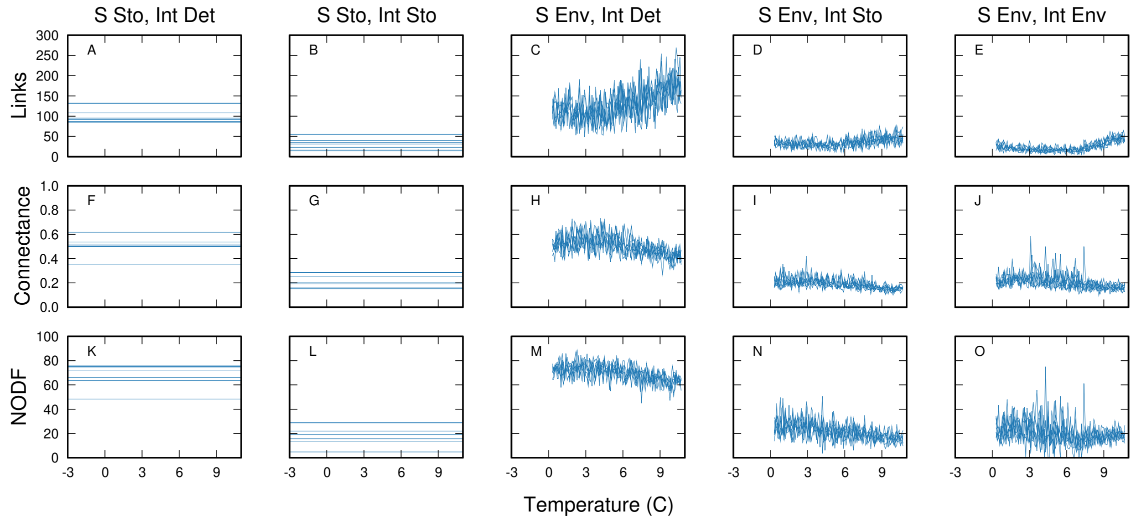


**Figure S11:** Properties of simulated plant-pollinator networks over the entire observed temperature range, 1996 networks shown. Networks were simulated for each 0.1 degree increment within the range shown.

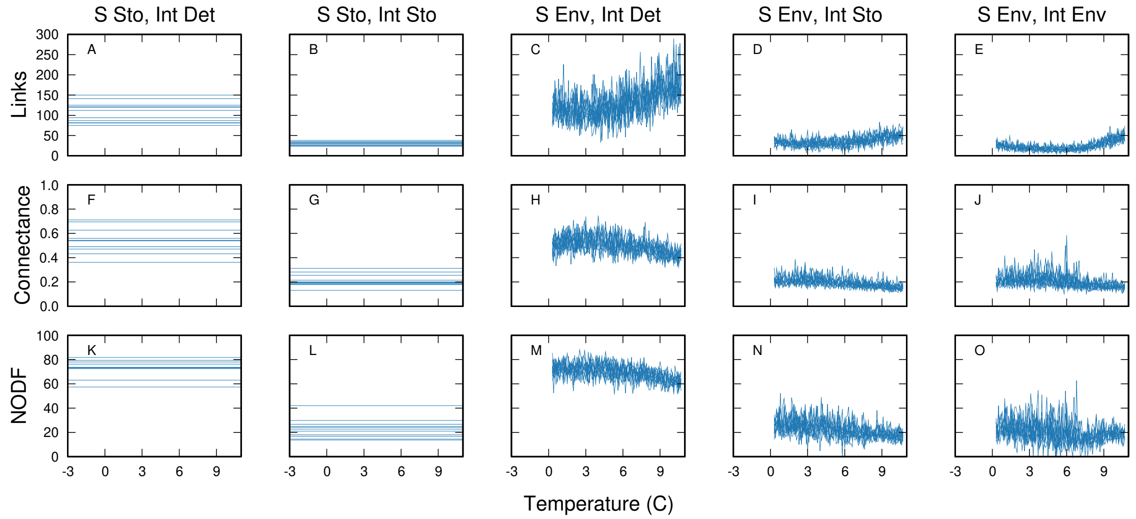


**Figure S12:** Properties of simulated plant-pollinator networks over the entire observed temperature range, 1997 networks shown. Networks were simulated for each 0.1 degree increment within the range shown.

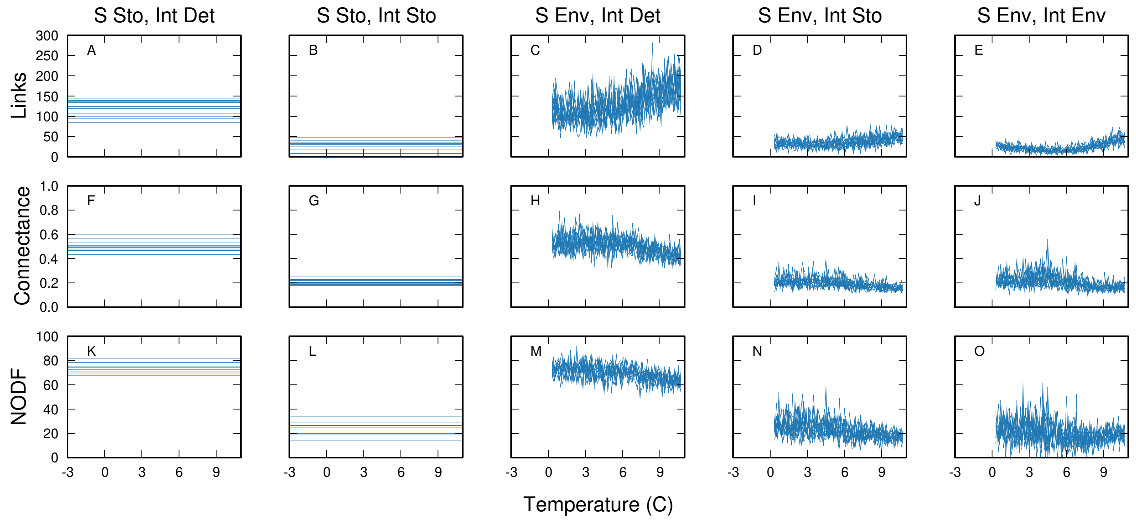


**Figure S13:** Properties of simulated plant-pollinator networks over the entire observed temperature range, 2010 networks shown. Networks were simulated for each 0.1 degree increment within the range shown.

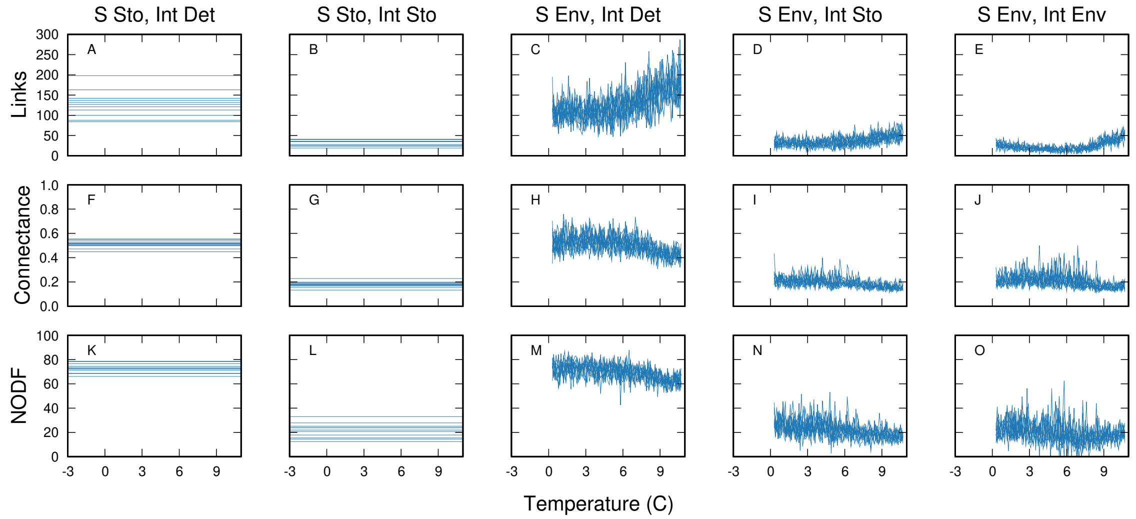


**Figure S14:** Properties of simulated plant-pollinator networks over the entire observed temperature range, 2011 networks shown. Networks were simulated for each 0.1 degree increment within the range shown.

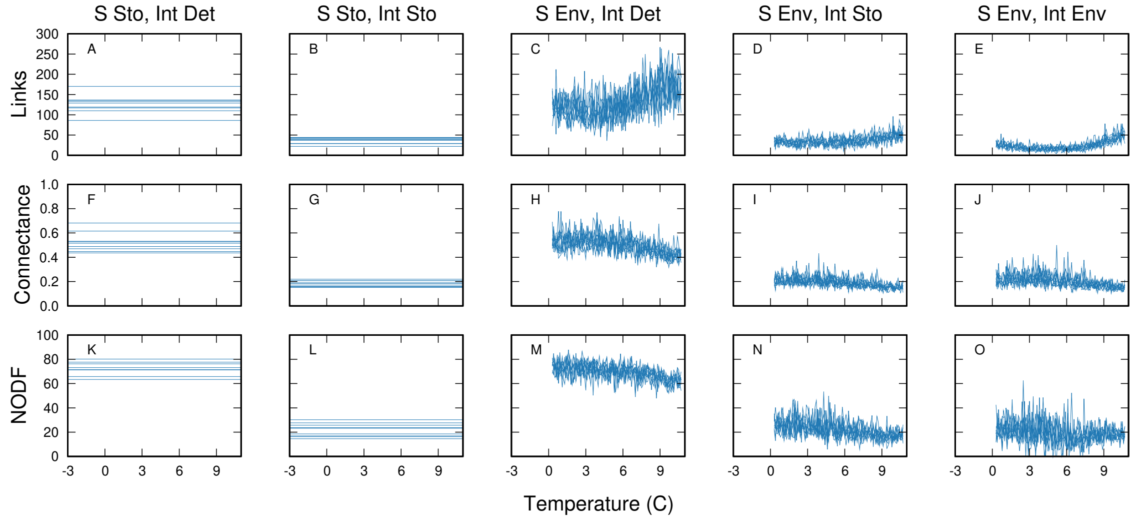


**Figure S15:** Properties of simulated plant-pollinator networks over the entire observed temperature range, 2016 networks shown. Networks were simulated for each 0.1 degree increment within the range shown.

### S5: Alternate display of observed and simulated network properties

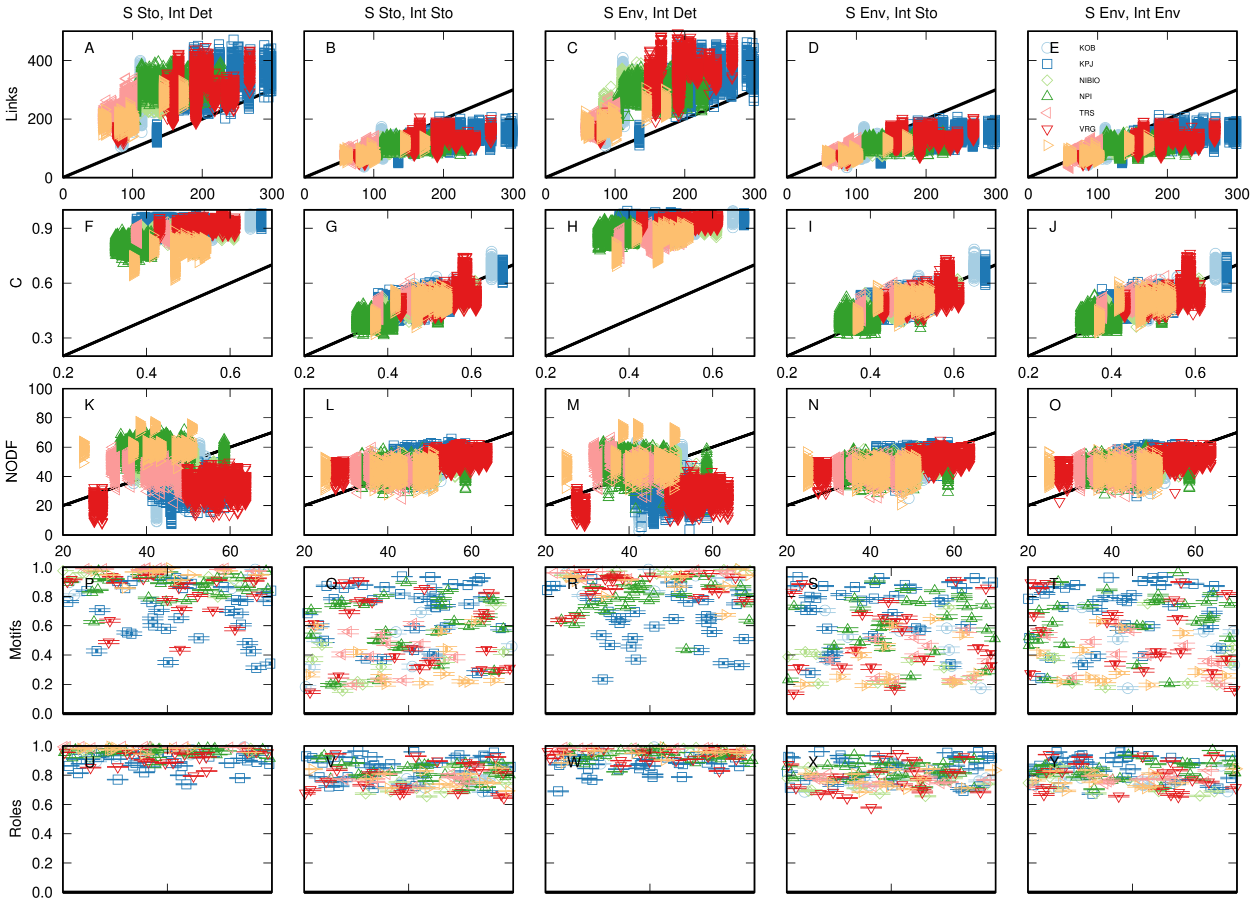


**Figure S16:** In the plant-fungus networks, the match between observed network properties (x-axis) and simulated network properties (y-axis) varied depending on the simulation strategy used, except for network motif profiles and species motif roles, where Bray-Curtis dissimilarity between observed and simulated networks (y-axis) was generally high regardless of simulation approach. Networks are randomly distributed across the x-axis for motif and role panels. Note that assuming that interactions occur deterministically can result in unrealistic network properties, even when species occurrence is modelled based on environmental niches.

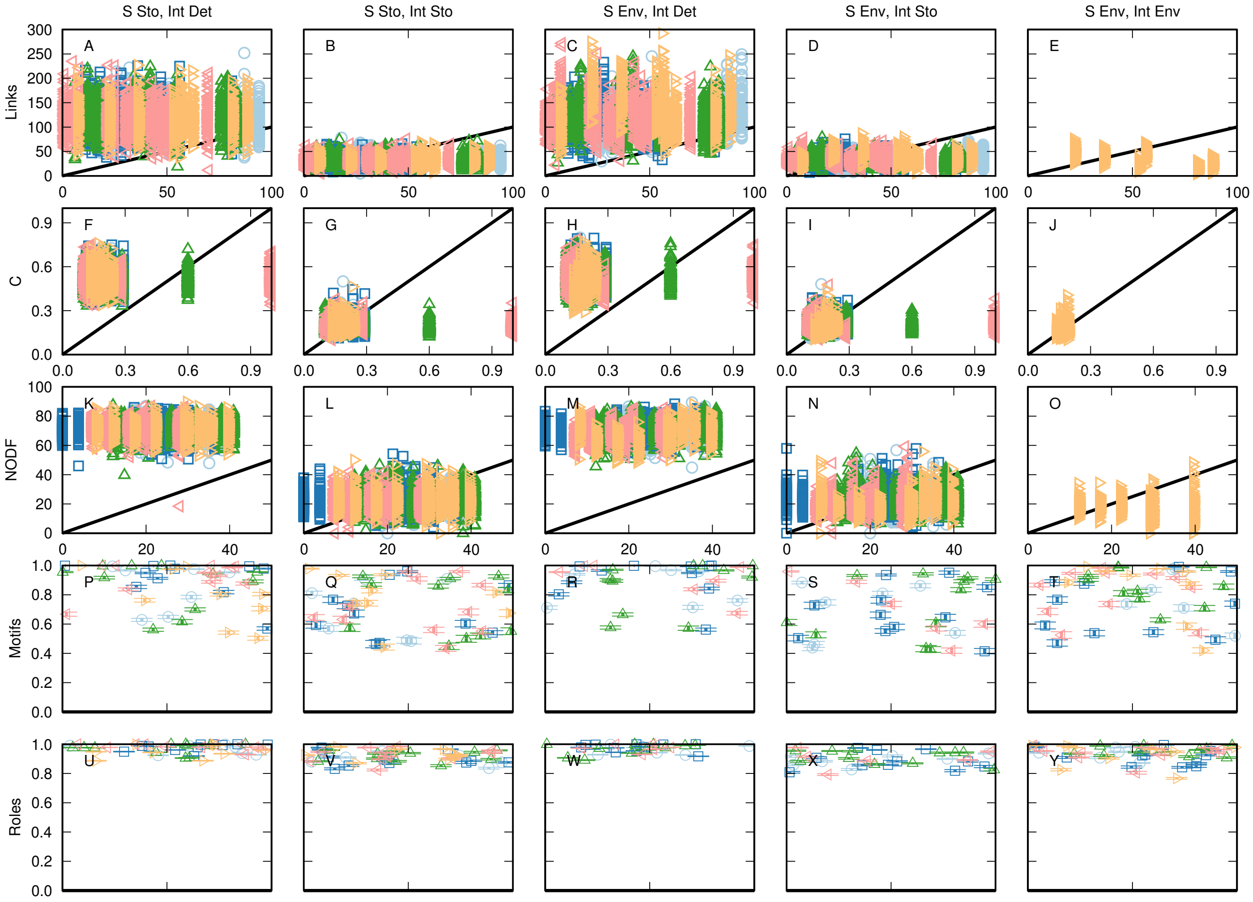


**Figure S17:** In the weekly plant-pollinator networks, the match between observed network properties (x-axis) and simulated network properties (y-axis) varied depending on the simulation strategy used, except for network motif profiles and species motif roles, where Bray-Curtis dissimilarity between observed and simulated networks (y-axis) was generally high regardless of simulation approach. Networks are randomly distributed across the x-axis for motif and role panels. Note that assuming that interactions occur deterministically can result in unrealistic network properties, even when species occurrence is modelled based on environmental niches.
